# Natural Antimicrobial Peptides Self-assemble as α/β Chameleon Amyloids

**DOI:** 10.1101/2022.06.23.497336

**Authors:** Peleg Ragonis-Bachar, Bader Rayan, Eilon Barnea, Yizhaq Engelberg, Alexander Upcher, Meytal Landau

**Affiliations:** Department of Biology, Technion-Israel Institute of Technology, Haifa 3200003, Israel; Ilse Katz Institute for Nanoscale Science and Technology, Ben Gurion University of the Negev, Beer-Sheva 84105, Israel; European Molecular Biology Laboratory (EMBL) and Centre for Structural Systems Biology, Hamburg 22607, Germany

**Keywords:** Antimicrobial peptides, amyloid, cross-α/β, fibril, secondary structure switch

## Abstract

Amyloid protein fibrils and some antimicrobial peptides (AMPs) share biophysical and structural properties. This observation suggests that ordered self-assembly can act as an AMP-regulating mechanism, and, vice versa, that human amyloids play a role in host defense against pathogens, as opposed to their common association with neurodegenerative and systemic diseases. Based on previous structural information on toxic amyloid peptides, we developed a sequence-based bioinformatics platform and, led by its predictions, experimentally identified 14 fibril-forming AMPs (ffAMPs) from living organisms, which demonstrated cross-β and cross-α amyloid properties. The results support the amyloid-antimicrobial link. The high prevalence of ffAMPs produced by amphibians and marine creatures among other species suggests that they confer unique advantageous properties in distinctive environments, potentially providing stability and adherence properties. Most of the newly identified 14 ffAMPs showed lipid-induced and/or time-dependent secondary structure transitions in the fibril form, indicating structural and functional cross-α/β chameleons. Specifically, ffAMP cytotoxicity against human cells correlated with inherent or lipid-induced α-helical fibril structure. The findings raise hypotheses about the role of fibril secondary structure switching in regulation of processes, such as the transition between a stable storage conformation and an active state with toxicity against specific cell types.

## INTRODUCTION

The emergence of resistant and aggressive microbial infections calls for novel and effective drugs. Antimicrobial peptides (AMPs) are naturally produced by many organisms as the first line of defense against pathogens and for modulation of the immune system^1^. AMPs have been extensively studied due to their potential for broad-spectrum and rapid activity, and lower likelihood to induce resistance as compared to conventional small-molecule antibiotics^2^. Yet, their relatively low efficacy and bioavailability, and lack of chemical stability curbed their development into therapeutic agents^2^.

Several AMPs assemble into well-ordered fibrils with amyloidogenic features such as amyloid-indicator dye binding and cross**-**β structures of tightly mated β-sheets^3–10^. Correspondingly, human amyloids associated with neurodegenerative and systemic diseases demonstrate antimicrobial activity, including amyloid-β (Aβ) and tau involved in Alzheimer’s disease, α-synuclein involved in Parkinson’s disease, serum amyloid A and beta2-microglobulin associated with systemic amyloidosis, human islet amyloid polypeptide involved in diabetes, and the human prion protein^11–25^. While these human amyloids share cross**-**β features, another amyloid form, named cross-α, is composed entirely of amphipathic α-helices that stack perpendicular to the fibril axis into mated ‘sheets’, reminiscent of the cross-β arrangement, and correspondingly, share amyloid-indicator dye binding^8,26,27^.

The cross-α fibril form, which was suggested to serve as a toxic functional amyloid form^8,26,27^, was revealed in the crystal structure of the cytotoxic phenol-soluble modulin α3 (PSMα3) peptide secreted by the pathogenic *Staphylococcus aureus* bacterium^26,27^, and later in the crystal structure of the uperin 3.5 AMP secreted by *Uperoleia mjobergii* (Mjoberg’s toadlet)^8^. The amyloid-forming uperin 3.5^8,28–30^ showed a secondary structure switch between cross-α and cross-β fibrils, depending on environmental conditions. In particular, exposure to lipids and surfactants drove a transition to an α-helical state^8,31^. We recently determined the high-resolution structure of the cross-β form of uperin 3.5 by cryogenic electron microscopy (cryo-EM)^32^. Specifically, uperin 3.5, dissolved in double-distilled water (ddH2O), displayed polymorphic fibrils with different arrangements based on a protofilament of two mated β-sheets, which is the hallmark of typical amyloids. The most abundant polymorph, determined at high resolution, showed a 3-blade symmetrical propeller of nine peptides per fibril layer including tight β-sheet interfaces^32^. The findings overall exposed the substantial polymorphism of the uperin 3.5 sequence, and, together, with the cross-α crystal structure, indicated the ability of fibril formation to drive different secondary structures depending on the environment and potential physiological needs^8,31,32^.

In light of the above findings, we hypothesized that fibril-forming AMPs (ffAMPs) displaying a secondary structure switch, might have adopted a regulation mechanism of switching between fibril polymorphs used for storage and adherence, versus for toxicity against different cell types induced by encountering cell membranes. To this end, this work developed a sequence-based bioinformatics platform to identify ffAMPs based on predictions of amphipathic α-helical together with β secondary structure propensities, herein termed chameleon sequences. In total, 14 AMPs from living organisms displaying fibril-forming capabilities and amyloidogenic properties were identified. Fibril helical secondary structure and lipid-induced switch correlated with cytotoxic activities. The presented findings reinforce the connection between antimicrobial activity and amyloidogenic features, which, in turn, supports the hypothesis that human amyloids associated with disease play a physiological role as part of the host defense system^33^. In addition, this link advocates the functional utilization of amyloid stability and adherence properties^34,35^ to enhance antimicrobial defenses across kingdoms of life. A more comprehensive understanding of the molecular properties of antimicrobial amyloids will promote the design of therapeutics and coating agents in the fight against pathogens.

### Experimental Section

#### Computational predictions

The ‘collection of anti-microbial peptides’ (CAMP_R3_) database of AMPs from eukaryotic and prokaryotic organisms^36^ was used as a source to screen for ffAMPs. Two independent prediction routes were taken: one to identify short helical AMPs, with prediction for helicity conducted with the Jpred server^37^, and one to identify chameleon AMPs, namely sequences that show propensities for both α-helical and β-structures, conducted with the Tango server^38^. For initial evaluation of secondary structure predictions and optimization of threshold values, AMPs were divided into two groups, i.e., with unknown and known 3D structures.

Some adaptations were required for the Jpred and Tango predictions. Specifically, for Jpred, amidation of C- and N-termini was removed (if existed), and peptides smaller than 20 amino acids (aa) were duplicated 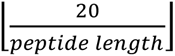 times to obtain a minimum length of 20 aa. For Tango predictions, the pH, temperature, and ionic strength were adjusted to physiological conditions. In case of undefined residues, they were substituted to known amino acids as follow: “X” was substituted to alanine; “Z” was substituted to glutamate; “B” was substituted to aspartate; and “U” was substituted to cysteine. None of the 14 ffAMPs reported here had these undefined residues.

Thresholds of the search were set according to the following parameters: The minimal length of consecutive residues predicted to have the same secondary structure was set to either 5 or 6 residues. The minimal gap length, i.e., the minimal number of sequential residues with no reliable secondary structure prediction located between residues with a reliable secondary structure prediction, was set to 3 residues. The minimal reliability score of α-helical prediction by Jpred was set to either 7 or 7.5. The minimal reliability score of α-helical predictions by Tango was either unlimited or set to 0.3, and the minimal reliability score of β prediction by Tango was either unlimited or set to 0.6. The maximal difference between α-helix and β prediction scores by Tango was either unlimited or set to 3. The minimal hydrophobic moment (µH) of the residues predicted to assume a helical secondary structure was set to either 0.8 or 0.7 for Jpred and either 0.8 or 0.61 for Tango. All threshold values used described in supplementary Table 1. The experimental assessment performed here indicated that the set µH threshold was too high. The reliability score of a peptide (in Jpred or Tango) were calculated as the average reliability scores of the segments predicted to have secondary structure. Peptides were predicted to have secondary structure if they passed all the thresholds, and their average or median reliability score passed the minimal threshold.

#### Materials

Peptides (>98% purity) were all purchased from GL Biochem (Shanghai) Ltd. A list of all peptide sequences, identification codes and properties, is presented in supplementary Tables 2,4&5. Dimethyl sulfoxide (DMSO), chloroform, methanol, 1,1,1,3,3,3-hexafluoro-2-propanol (HFIP), thioflavin T (ThT), deuterium oxide (D2O), and uranyl acetate, were purchased from Sigma-Aldrich. Anhydrous sodium phosphate, monobasic NaH2PO4 was purchased from Merck. Ultra-pure water was purchased from Biological Industries (Israel). Luria-Bertani (LB) bacterial medium was purchased from Difco-BD, USA. Lipids, including 1,2-dioleoyl-sn-glycero3-phosphoethanolamine (DOPE),1,2-dioleoyl-sn-glycero-3-phospho- (1’-racglycerol) (DOPG), 1,2-dioleoyl-sn-glycero-3-phosphocholine (DOPC), cholesterol (Chol) and sphingomyelin (SM) were purchased from Avanti Polar Lipids, Inc. (USA).

#### Peptide pre-treatment

All AMPs were dissolved in HFIP to a concentration of 1 mg/ml, followed by a 10 min bath-sonication, at room temperature. The organic solvent was evaporated using a mini rotational vacuum concentrator (Christ, Germany) at 1,000 rpm for 2 h, at room temperature. Treated peptides were aliquoted and stored at -20 °C.

#### Bacterial strains and culture media

*Micrococcus luteus* (an environmental isolate) was a kind gift from Prof. Charles Greenblatt from the Hebrew University of Jerusalem, Israel. An inoculum was grown in LB medium at 30 °C, with 220 rpm shaking, overnight.

#### Antibacterial activity in solution

Minimal inhibitory concentration (MIC) was defined as the minimal concentration in which bacterial growth was reduced to 20% or less over 24 h. Bacterial cultures were grown overnight, as described above, and diluted 1,000-fold in fresh medium until OD_595 nm_ reached ∼0.4. Peptides were dissolved to a concentration of 1 mM in ultra-pure water and diluted to 500 μM in LB. Two-fold serial dilutions in LB of the tested peptides (250 μM to 0.5 μM) were performed in a sterile 96-well plate. Wells containing everything but the peptide served as controls. Bacterial growth was determined by measuring OD_595 nm_ during the 24 h incubation (30°C, 220 rpm shaking). The experiment was performed in triplicates. All triplicate values were averaged, appropriate blanks were subtracted, and MIC was determined according to the definition above. The experiment was repeated 3 times on different days.

#### Lactate dehydrogenase (LDH) release cytotoxicity assay

Human lymphoblast T2 cells (174 x CEM.T2) (ATCC® CRL-1992™) were routinely cultured in RPMI 1640 medium (Sigma, Israel) with L-glutamine, supplemented with penicillin (100 U/ml), streptomycin (0.1 mg/ml), and 10% heat-inactivated fetal calf serum (Biological Industries, Israel), at 37°C and 5% CO2. ffAMPs were dissolved to 1 mM in ultra-pure water, sonicated for 10 min, and diluted in RPMI 1640 medium with L-glutamine supplemented with penicillin (100 U/ml), streptomycin (0.1 mg/ml), and 0.5% heat-inactivated fetal calf serum (assay medium) to 160 μM. Serial two-fold dilutions in assay medium were performed, and 50 μl of each dilution were pipetted into a 96-well plate in triplicates and incubated for 15 min, at room temperature. The cells were washed and resuspended in assay medium to 0.15 × 10^6^ cells/ml, 50 μl of which were added to the diluted peptides and incubated for 30 min at 37 °C, 5% CO2. Cell lysis was quantified using the LDH release colorimetric assay according to the manufacturer’s instructions, including all recommended controls (LDH; Cytotoxicity Detection Kit Plus, Roche Applied Science, Germany). Cell-free assay medium was measured as background. Cells subjected to the same experimental conditions apart from peptide addition, served as a control to account for spontaneous LDH release. Cells subjected to the same experimental conditions apart from peptide addition and treated with manufacturer-supplied lysis buffer, served as a positive control to determine maximum LDH release. Absorbance at 490 and 690 nm was measured by a plate reader (FLUOstar Omega, BMG Labtech, Germany). Absorbance values at 690 nm were subtracted from absorbance values at 490 nm, average absorbance values of triplicate samples and controls were calculated, background was subtracted, and the 50% lethal concentration (LC50) values were determined. The experiment was repeated three times on different days.

#### Thioflavin T fluorescence fibrillation kinetics assay

ThT is a widely used “gold standard” stain for identifying and exploring amyloid fibril formation kinetics, both in vivo and in vitro. Fibrillation curves in the presence of ThT commonly show a lag time for the nucleation step, followed by rapid aggregation. All AMPs were dissolved in HFIP to a concentration of 1 mg/ml, followed by 10 min bath-sonication, at room temperature. The organic solvent was evaporated using a mini-rotational vacuum concentrator (Christ, Germany) at 1,000 rpm for 2 h, at room temperature. Treated peptides were aliquoted and stored at -20 °C prior to use.

Prior to the experiment, aliquots were dissolved 1:1 DMSO:UPddw to 10 mM and immediately diluted to 100 μM in phosphate buffer, pH 7.5, containing filtered ThT diluted from stock made in UPddw. Final concentrations for each reaction were 100 µM peptide and 200 µM ThT. Blank solutions were also prepared for each reaction, containing all components aside from the peptides. The reaction was carried out in black 96-well flat-bottom plates (Greiner bio-one) covered with a thermal seal film (EXCEL scientific) and incubated in a plate reader (OMEGA) at a temperature of 37°C with 500 rpm shaking for 85 s, before each reading cycle, and up to 1000 cycles of 6 min each. Measurements were made in triplicates. Fluorescence was measured by excitation at 438±20 nm and emission at 490±20 nm over a period of about 100 h. All triplicate values were averaged, appropriate blanks were subtracted, and the resulting values were plotted against time. Calculated standard errors of the mean are presented as error bars. The entire experiment was repeated at least three times on different days.

#### Transmission electron microscopy (TEM)

The peptides were dissolved in UPddw to a concentration of 1 mM and incubated at 37 °C for 7 days. To image peptides in the presence of bacterial cells, *M. luteus* was grown for 24 h in LB. Approximately 1.5 × 10^9^ bacteria cells were washed three times with 10 mM potassium phosphate buffer, pH 7.4. Citropin-1.3 and temporin-1CEa were dissolved in this same buffer and added to the bacterial pellets, which were resuspended to a final peptide concentration of 8 µM and 40 µM, respectively. Samples were then incubated, at 30 °C, with 220 rpm shaking, for 4 h.

Samples (4–5 µl) were applied directly onto glow-discharged (easiGlow; Pelco, Clovis, CA, USA, 15 mA current; negative charge; 25 s time) 400 mesh copper grids, with a grid hole size of 42 µm, stabilized with Formvar/carbon (Ted Pella, Inc.). Samples of peptide alone were allowed to adhere for 60 s, and samples with *M. luteus* were allowed to adhere for 45 s. Samples were then stained with 1% uranyl acetate solution (Electron Microscopy Science, 22400-1) for 60 s (peptide alone) or 45s (peptide with *M. luteus*), before being blotted with Whatman filter paper. Samples of peptides alone were examined with a ThermoFisher Scientific (FEI) Tecnai T12 G2 electron microscope, at an accelerating voltage of 120 kV, equipped with Gatan 794 MultiScan CCD camera, or with ThermoFisher Scientific (FEI) Talos F200C transmission electron microscope operating at 200 kV, equipped with Ceta 16M CMOS camera. Samples of peptides incubated with *M. luteus* were examined with a FEI Tecnai G2 T20 electron microscope, at an accelerating voltage of 200 kV.

#### Small unilamellar vesicle (SUV) preparation

Lipids were dissolved to 150 mM stock solutions in 1:1 methanol:chloroform and then mixed in the following combinations and ratios:1,2-dioleoyl-sn-glycero-3-phosphoethanolamine (DOPE) and 1,2-dioleoyl-sn-glycero-3-phospho-(1′-rac-glycerol) (DOPG) at a 1:1 molar ratio, mimicking a Gram-positive bacterial membrane, or 1,2-dioleoyl-sn-glycero-3-phosphocholine (DOPC), cholesterol(Chol) and sphingomyelin (SM) at a 0.67:0.25:0.08 molar ratio, mimicking an eukaryotic membrane. The solvent was then evaporated under vacuum, to complete dryness. The lipid film was rehydrated in 10 mM potassium phosphate buffer, pH 7.5, for the CD measurements and in 10 mM anhydrous sodium phosphate, monobasic, NaH2PO4 in D2O buffer, which was titrated using HCl to pH 7.5, for the FTIR measurements. The samples were vortexed and incubated for several hours to overnight. The final lipid concentration was 10 mM. The lipid suspension was sonicated, on ice, using a tip sonicator (SONICS, USA), at 20 % amplitude, for 5 min, with 10 s pulses, to form SUVs. Prior to use, the solution was incubated for 1 h at room temperature.

#### Solution circular dichroism (CD) spectroscopy

Immediately prior to CD experiments, peptide samples were dissolved in ultra-pure water to 10 mM and diluted to 1 mM in 10 mM potassium phosphate. The signal from blank solutions of either buffer alone or of buffer with SUVs composed of either DOPE:DOPG (bacterial membrane model) or DOPC:Chol:SM (eukaryotic membrane model), were recorded before the addition of the peptide sample to the cuvette. The peptides were diluted to a concentration of 0.2 mM, in the presence or absence of 0.6 mM SUVs to obtain a peptide:lipid ratio of 1:3. Far-UV CD spectra were recorded with an Applied Photophysics PiStar CD spectrometer (Surrey, UK), using a 1 mm path-length quartz cell (Starna Scientific, UK), at a temperature of 25 °C. Changes in ellipticity were monitored from 260 nm to 180 nm, at 1 nm steps and a bandwidth of 1 nm. The measurements shown are an average of five scans for each sample, captured at a scan rate of 1 sec per point, with appropriate blanks subtracted. Due to peptide aggregation upon dilution into the tested buffer, which significantly lowered the soluble peptide concentration, the concentration of each sample at real time was experimentally determined using UV absorbance, measured in parallel to CD spectra measurements, taken by the same machine using the Beer-Lambert equation at 205 nm: A_205nm_ = ε205×l×C , where ε205nm (M-1 x cm-1) is a sequence-specific extinction coefficient calculated using the webserver (http://spin.niddk.nih.gov/clore/). The pathlength ‘l’ was 1 mm. A_205nm_ was determined by averaging the blank corrected absorbance values at 205 nm from all four scans. The molar ellipticity per residue (Ɵ, in mdeg*cm2 *dmol-1*residu-1) was derived from the raw data (δ, in mdeg), using the following formula: Ɵ = δ / (L*C*N), where L is the path length of the cuvette, C is the real-time calculated molar concentration of the peptide solution, and N is the number of residues of the peptide. Deconvolution of the resulting CD spectra to determine the secondary structure composition was done using the BestSel webserver (http://bestsel.elte.hu/).

#### Attenuated total internal reflections (ATR) Fourier transform infrared (FTIR) spectroscopy

All peptides were dissolved in HFIP to a concentration of 1 mg/ml, then frozen in liquid nitrogen and lyophilized for several hours to complete dryness. Next, the peptides were dissolved to 1 mg/ml in 5 mM hydrochloric acid (HCl) and sonicated in a bath sonicator for 5 min, at room temperature. The peptide solution was then frozen in liquid nitrogen and lyophilized overnight to complete dryness. The procedure was repeated twice to completely remove trifluoroacetic acid (TFA) residues, as TFA has a strong FTIR signal at the amide I’ region of the spectra. Finally, the peptides were dissolved in D2O to 1 mg/ml, frozen in liquid nitrogen and lyophilized overnight to complete dryness. The procedure was repeated twice. For the time-dependent FTIR experiments, the dry peptides were dissolved in D2O to a concentration of 10 mM and samples were measured after 2 h, 24 h, and 72 h of incubation. To test the effect of the lipids, the dry peptides were dissolved to a final concentration of 10 mM immediately prior to measurements in the presence or absence of DOPE:DOPG and DOPC:Chol:SM SUVs, in the same 10mM sodium phosphate buffer used to prepare the SUVs, and incubated for 3 h.

Samples (5 μl) were spread on the surface of the ATR module and allowed to dry. Absorption spectra were recorded on the dry samples using a Tensor 27 FTIR spectrometer (Bruker Optics). Measurements were performed in the wavelength range of 400-4000cm^-1^ in 2 cm^-1^ steps and averaged over 32 scans. Absorbance of background (Air) and blank (D2O, Buffer, DOPE:DOPG or DOPC:Chol:SM SUVs in D2O) were measured and showed a negligible signal. The peaks were selected by the instrument based on the second derivative using OPUS software.

#### Fiber X-ray diffraction

The peptides were dissolved in double-distilled water to 10 mM and incubated between 2 h to 7 days, as indicated. Droplets (2 μl) were placed between two sealed-end glass capillaries and incubated until complete dryness (up to 1 h) at room temperature. X-ray diffraction of the fibrils was collected at the micro-focus beamline P14, operated by EMBL Hamburg, at the PETRA III storage ring (DESY, Hamburg, Germany). The experiment was repeated twice on different days.

## RESULTS

### Prediction of α-helical and chameleon fibril-forming AMPs (ffAMPs)

The recently solved structures of toxic fibril-forming peptides that revealed helical fibrils^8,26,27,39–42^, especially of PSMα3 exposing the amyloid cross-α features^26,27,42^, and of uperin 3.5 featuring the chameleon cross-α/β switch^8,32^, inspired the search for additional ffAMPs based on specific sequence properties. The solved structures contained amphipathic helices that assembled via a hydrophobic core. In the cross-α form, the helices assembled via mated sheets that were glued together by an extensive hydrophobic core along the fibril axis. We therefore screened for AMPs which are short peptides (<100 amino acids), have high helical propensity or have propensities for both secondary structures, and a large hydrophobic moment (µH) for the helical regions.

The Collection of Anti-Microbial Peptides (CAMP_R3_) database^36^ was computationally screened for AMPs based on predictions of length, secondary structure and µH. Two prediction routes were pursued: identification of short helical AMPs, with helicity predicted using the Jpred server^37^, and identification of chameleon AMPs, namely sequences that show propensities for both α-helical and β-structures, executed using the Tango server that is designed to identify β-rich amyloidogenic sequences^38^. Three iterations were conducted testing several threshold values each, and predictions were compared to secondary structures of AMPs with known three-dimensional (3D) structures and to uperin 3.5 to refine parameters. The parameters and thresholds are outlined in the Method section, and the numeric results of the predictions with three different sets of thresholds are described in Supplementary Table 1.

After the third iteration, 26 naturally produced AMPs were selected and tested for fibril formation. Fourteen of the AMPs formed fibrils (Figure 1) and were further characterized. Sequences and references are provided in Supplementary Table 2, and physico-chemical properties along with predictions (helical and/or chameleon) are provided in Supplementary Table 3. Of note, fibril formation was also observed for some of our predicted AMPs which originated from designed synthetic sequences. However, the current work focused on sequences produced by live organisms.

**Figure 1.**
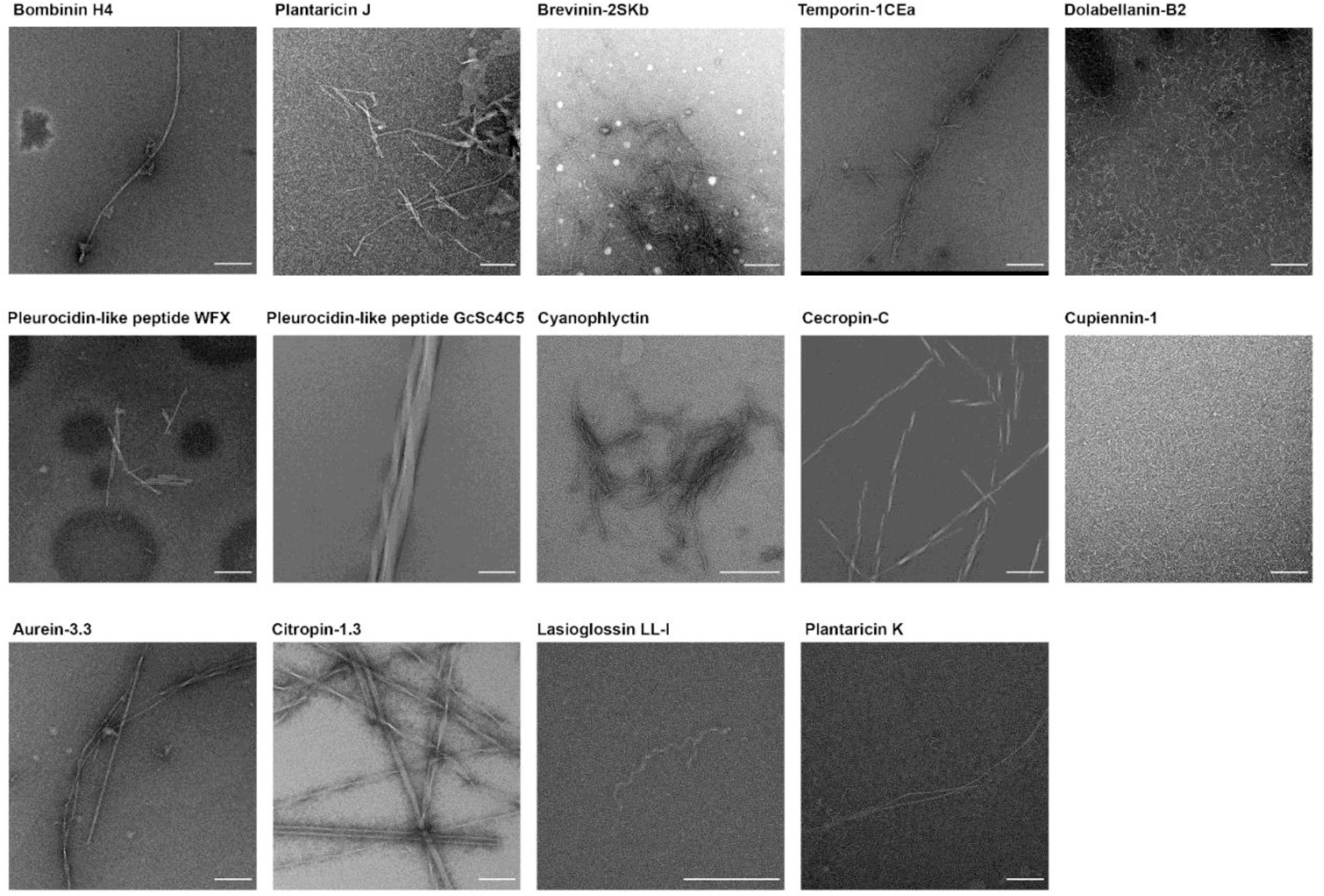
Fibril formation of ffAMPs visualized by transmission electron micrographs. Transmission electron micrographs of 1 mM fibril-forming antimicrobial peptides (ffAMPs) showing different fibril morphologies. All the peptides were incubated in ddH2O for one week. All scale bars represent 200 nm.

The 14 ffAMPs included cyanophlyctin, citropin-1.3, brevinin-2SKb, temporin-1CEa, aurein-3.3, and bombinin H4, all produced by amphibians, pleurocidin-like peptide GcSc4C5 and pleurocidin-like peptide WFX produced by fish, dolabellanin-B2 produced by a sea slug, cupiennin-1 produced by a spider, lasioglossin LL-I produced by a bee, cecropin-C produced by mosquitoes, and plantaricin-J and plantaricin-K produced by a bacterium. The producing organisms, and the CAMP_R3_ database identification codes^36^ are specified in Supplementary Table 4. In addition, Supplementary Table 5 provides database identification information on the ffAMPs, if available, including from Uniprot, GenBank of the National Center of Biotechnology (NCBI), and the protein data bank (PDB). Of note, cupiennin-1, brevinin-2SKb, plantaricin-K and bombinin H4 did not pass the thresholds for the hydrophobic moment set in Supplementary Table 1 and were studied as lower-threshold examples, which nevertheless showed fibril formation.

### The 14 ffAMPs display various fibril morphologies and amyloid-indicator dye binding abilities

The ffAMPs displayed fibrils with different morphologies visualized by transmission electron microscopy (TEM), (Figure 1). Five out of the 14 ffAMPs, including temporin-1CEa, bombinin H4, pleurocidin-like peptide GcSc4C5, citropin-1.3 and plantaricin-K, induced fluorescence by the binding of the amyloid-indicator dye thioflavin-T (ThT) (Supplementary Figure 1).

### Antimicrobial activity and cytotoxicity of the ffAMPs

The 14 ffAMPs were previously reported to have antimicrobial activity against different species. The reporting citation and the antimicrobial database reference numbers are provided in Supplementary Tables 2&4. Here, to compare activity levels against a specific microorganism we tested the antibacterial activity of the ffAMPs against *Micrococcus luteus* using the broth-dilution assay, by determining the minimum inhibitory concentration (MIC) of the freshly dissolved peptides. *M. luteus* was selected due to its low pathogenicity to humans and robust growth in conventional media. All but two ffAMPs proved active against *M. luteus*; pleurocidin-like peptide WFX and dolabellanin-B2 showed no antimicrobial effect at concentrations up to 250 μM (Table 1). Citropin-1.3 and temporin-1CEa incubated in the presence of *M. luteus* at a concentration around their MIC levels, formed fibrils around bacterial cells (Figure 2), suggesting fibril involvement in the antibacterial activity. The cytotoxicity of the ffAMPs was also measured against the immune-related T2 cells, and eight out of the 14 ffAMPs showed cytotoxic effects within the tested concentration range (Table 1).

**Figure 2.**
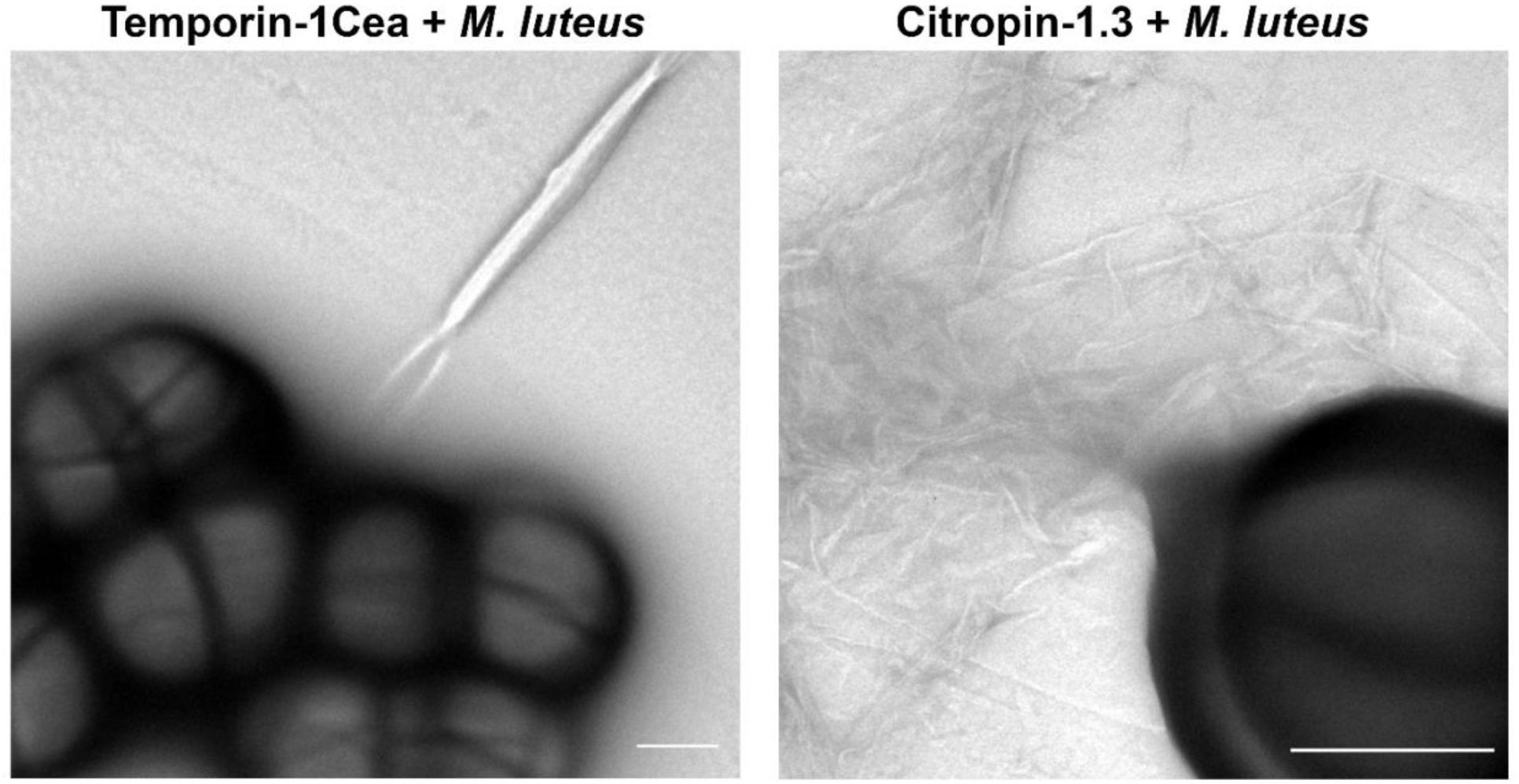
Temporin-1Cea and citropin-1.3 form fibrils around bacterial cells. Transmission electron micrographs of 40 µM temporin-1Cea (close to its MIC of 31 µM) and 8 µM citropin-1.3 (close to its MIC of 6.6 µM) incubated with *M. luteus* for 4 h. Both scale bars represent 500 nm.

**Table 1.**
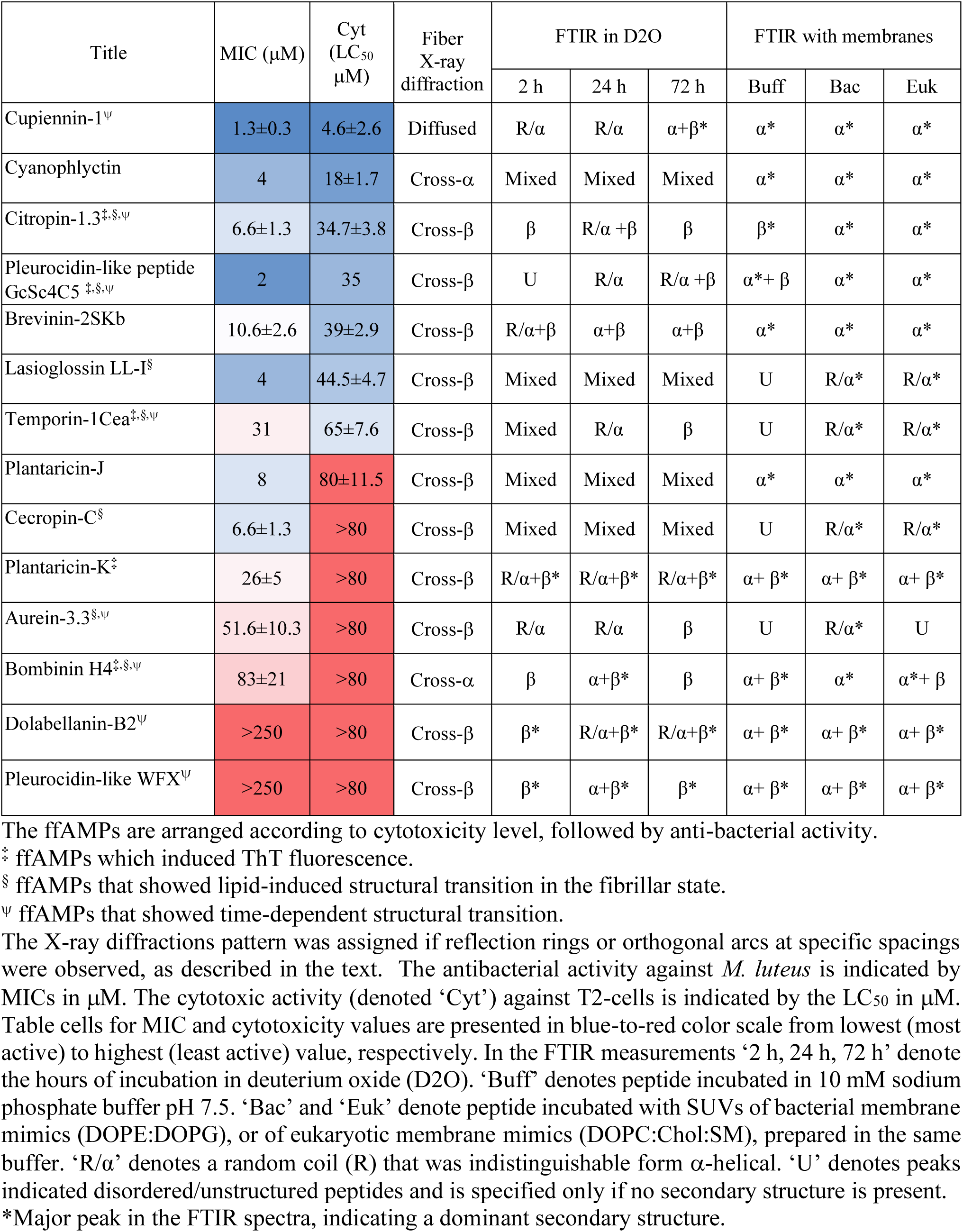
Toxic activities and secondary structures in the fibril form of the identified ffAMPs.

### The ffAMPs show amyloid X-ray fiber diffraction patterns

X-ray fiber diffraction analyses were performed to determine the presence of ordered structures of continuous sheets. A typical cross-β signature displays orthogonal reflection arcs at 4.6-4.8 Å spacing, corresponding to the distance between β-strand along the β-sheets (axial packing), and at 8-12 Å spacing, corresponding to the distance between pairs of β-sheets (lateral packing), with the latter reflection often being more diffuse^43^. The X-ray fiber diffraction of the cross-α structure of the *S. aureus* PSMα3 fibril displays a reflection arc at 10.5 Å spacing with orthogonal reflection arcs at 12 Å and 25 Å^26^. According to the PSMα3 crystal structure, the 10.5 Å and 12 Å reflections correspond to the spacing between the α-helices along each sheet (axial packing) and the inter-sheet distance (lateral packing), respectively^27^. The reflection at 25 Å may be attributable to the length of the helix, also being orthogonal to the distance between helices along the sheet, or to the distance between pairs of sheets, perpendicular to the fibril axis.

The X-ray fiber diffraction of 11 out of the 14 ffAMPs showed a cross-β signature with orthogonal reflection arcs or rings at 4.6-4.7 Å and 8-12 Å spacing. The arcs indicate higher order in the fibril alignment and/or arrangement compared to rings. Cross-β-forming temporin-1CEa and pleurocidin-like WFX are displayed in Figure 3; the remaining ffAMPs are displayed in Supplementary Figures 2&3. Cupiennin-1 showed diffused rings with 3.8-4.7 Å and 8-11 Å reflections, which might be attributable to a β-rich fibril, yet the lack of a sharp reflection at around 4.7 Å prevented us from classifying cupiennin-1 as cross-β in Table 1. Bombinin H4 and cyanophlyctin showed fiber diffraction that resembled the cross-α pattern of PSMα3^26^ (Figure 3), as detailed below. Three ffAMPs displayed structural changes in the fiber diffraction pattern measured after different incubation times or on different days (Supplementary Figure 3), as detailed below in the individual description of ffAMPs.

**Figure 3.**
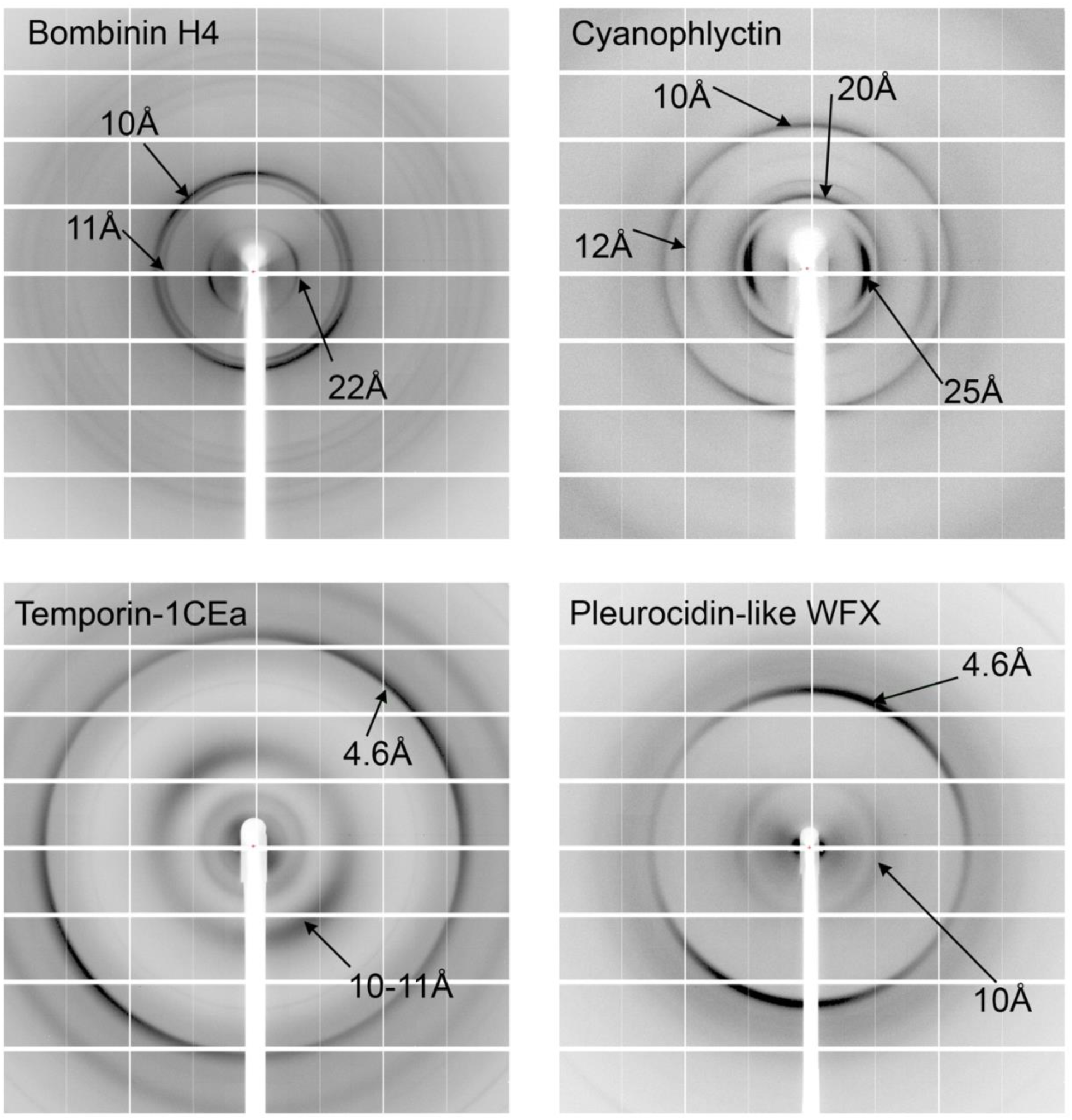
Cross-α/β X-ray fiber diffraction patterns of select ffAMPS. The X-ray fiber diffraction patterns of bombinin H4 (incubated for 6 h), cyanophyctin (incubated for 2 h), pleurocidin-like WFX (incubated for 2 h), and temporin-1Cea (incubated for 6 h). All ffAMPs were incubated at 10 mM in ddH2O. Major reflections are indicated.

### Secondary structures in soluble and fibrillar states and in the presence of lipids mimicking bacterial and eukaryotic membranes

While X-ray fiber diffraction denotes the presence of an amyloid arrangement, it cannot indicate the variety and abundance of fibril polymorphs. The indicated reflections can originate from a dominant or minor polymorph, while arrangements lacking a continuous structure with repeated spacings are likely to be undetectable by X-ray diffraction. Thus, circular dichroism (CD) spectroscopy and attenuated total internal reflection Fourier transform infrared (ATR-FTIR) spectroscopy which was shown to be useful for characterization of fibrillar architectures in amyloid proteins and peptides^44–48^, were used to assess the spectrum of secondary structure arrangements of the ffAMPs, in solution and in fibrils. The secondary structure of the ffAMPs was compared after an incubation in 10mM sodium phosphate buffer, pH 7.5, in the presence or absence of small unilamellar vesicles (SUVs) composed of either 1,2-dioleoyl-sn-glycero-3-phosphoethanolamine (DOPE) and 1,2-dioleoyl-sn-glycero-3-phospho-(1′-rac-glycerol) (DOPG) at a 1:1 molar ratio, mimicking a Gram-positive bacterial membrane ^49,50^, or 1,2-dioleoyl-sn-glycero-3-phosphocholine (DOPC), cholesterol (Chol) and sphingomyelin (SM) at a 0.67:0.25:0.08 molar ratio, mimicking an eukaryotic membrane ^51^.

In solutions free of lipids, most ffAMPs were unstructured (with typical minima at 197 nm in the CD spectra) except for aurein-3.3, which adopted an α-helical conformation (with typical minima near 208 and 222 nm), and dolabellanin-B2, which adopted a β-rich conformation (with typical minima near 218 nm) (Supplementary Figure 4 and Supplementary Table 6). Neither one showed secondary structure transition upon addition of lipids to the solution. In contrast, five other ffAMPs showed a lipid-induced structural transition toward an α-helical conformation in solution. Citropin-1.3 and temporin-1Cea transitioned from an unstructured state towards an α-helical conformation in the presence of SUVs, mimicking either bacterial or eukaryotic membranes (Supplementary Figure 4). For bombinin H4, brevinin-2SKb, and pleurocidin-like peptide GcSc4C5, only the presence of bacterial-, but not eukaryotic-, membrane mimics induced such changes in solution (Supplementary Figure 4), suggesting specificity in the interactions with different cell types.

For the ATR-FTIR measurement, in addition to comparison with and without the presence of SUVs, the ffAMPs secondary structure was also characterized over time, in order to observe time-dependent structural transitions in the fibrillar state. The results are presented in Figure 4 and Supplementary Figure 5, and summarized in Table 1. In the FTIR spectra, the secondary derivative of each curve was calculated and depicted to resolve and identify the major peaks contributing to the ATR-FTIR overlapping signal in this region. Peaks in the region of 1611–1630 cm^-1^ as well as ∼1685–1695 cm^-1^ are indicative of rigid cross-β fibrils. Peaks in the region of 1637–1645 cm^-1^ indicate disordered species, partially overlapping with peaks at 1630–1643 cm^-1^ correlating with small and disordered β-rich amyloid fibrils with absorbance typical of bent β-sheets in native proteins^48,52–54^. Peaks in the region of 1645–1662 cm^-1^ indicate the existence of α-helices^48,52–54^, but may overlap with random coil structures, especially for broad peaks^55,56^; therefore, both options are indicated (“R/α”) in Table 1. In case of a broad peak that covered both α, β and disordered ranges, the secondary structure is denoted as “mixed” in Table 1. Of note, a permissive range of peaks was used, in which we indicated helical structure according to observations made in a range of globular proteins^57^. We also considered that stacks of helices, as in the cross-α structure, might have a shifted peak compared to helices in solution. Overall, eight ffAMPs displayed a time-dependent secondary structure transition in the fibril form, and seven ffAMPs displayed a lipid-induced shift into a more α-helical content in the fibrillar state, as detailed below. In addition, incubation of the peptides in 10mM sodium phosphate buffer, pH 7.5, often reduced population dispersiveness and tended to strengthen the α-helical conformation compared to peptides dissolved and incubated in D2O (Figure 4, Supplementary Figure 5, and Table 1).

**Figure 4.**
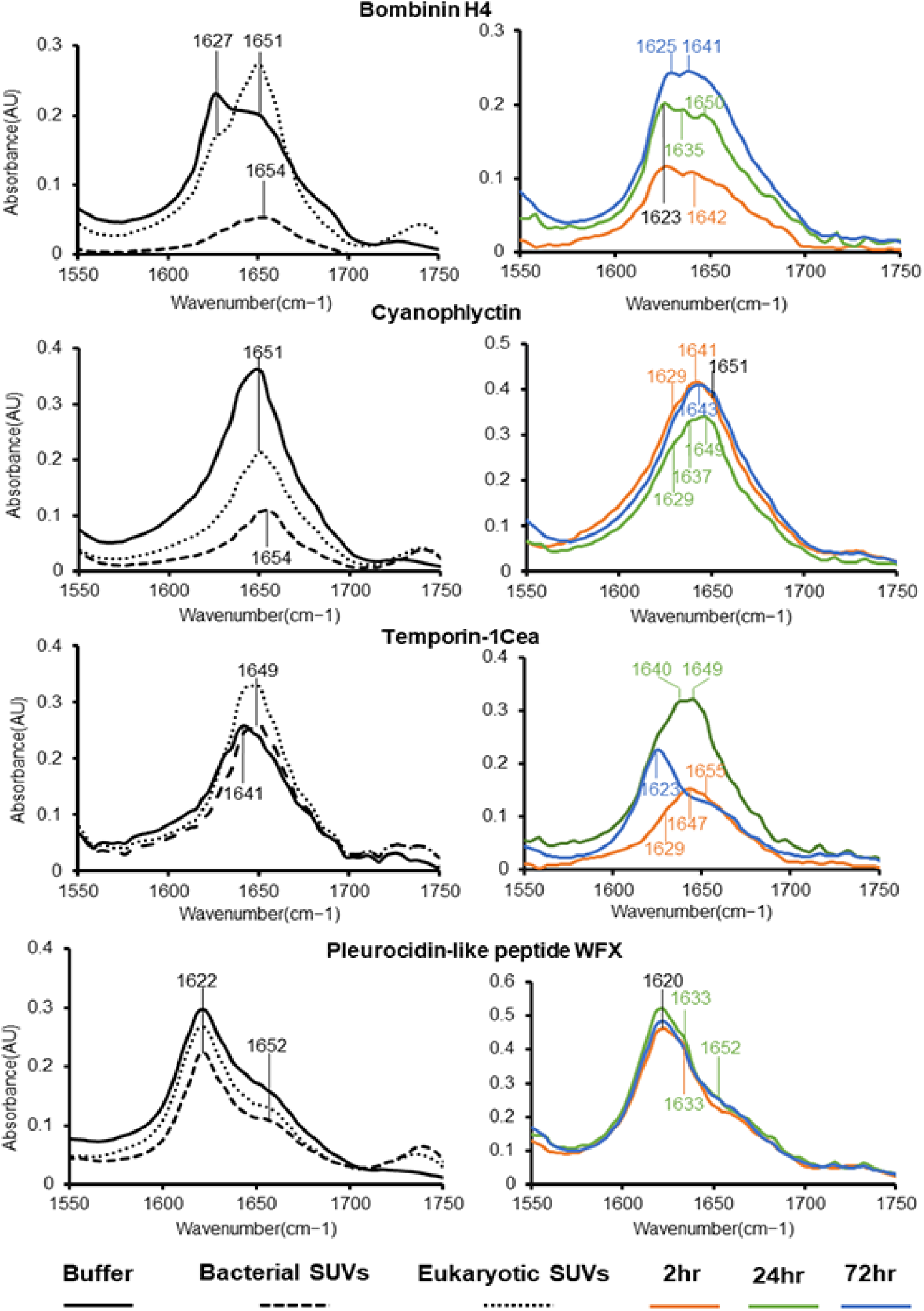
Secondary structure analyses of four representative ffAMPs in the fibrillar state. ATR-FTIR spectra of the amide I′ region of bombinin H4, cyanophlyctin, temporin-1Cea and plantaricin-K under various conditions and over time. Left: spectra are of the peptides incubated for several hours in 10 mM sodium phosphate buffer pH 7.5 (solid curve), or in the presence of SUVs composed of 1,2-dioleoyl-sn-glycero-3-phosphoethanolamine (DOPE) and 1,2-dioleoyl-sn-glycero-3-phospho-(1′-rac-glycerol) (DOPG), mimicking a Gram-positive bacterial membrane (dashed curve) or 1,2-dioleoyl-sn-glycero-3-phosphocholine (DOPC), cholesterol (Chol) and sphingomyelin (SM) mimicking a eukaryotic membrane (dotted curve). Right: spectra are of the peptides incubated in D2O for 2 h (orange curve), 24 h (green curve) or 72 h (blue curve). The peaks were determined based on the calculated second derivative provided by OPUS software.

### Biophysical properties of individual ffAMPs

#### Cupiennin-1

Cupiennin-1, originating from the spider venom of *Cupiennius salei*, shows potent antimicrobial, hemolytic, and insecticidal activity^58^. In the current analysis, it proved to be the most toxic peptide against both *M. luteus* (MIC of 1.3 μM) and T2-cells (LC_50_ of 4.6μM) (Table 1). Structurally, when incubated in D2O over time, cupiennin-1 showed a noticeable time-dependent shift from α-helical or random coil structures (peaks 1643-1651 cm^−1^) towards a predominantly cross-β content, with a peak at 1620 cm^−1^ after 72 h, along with a minor α-helical structure. The X-ray fiber diffraction of cupiennin-1, taken after several hours of incubation, showed a potential but not well-defined cross-β pattern (Supplementary Figure 2). In buffer with or without lipids, the α-helical fibrillar structure was dominant, with a relatively sharp peak in the FTIR spectra range of 1647-1649 cm^−1^ (Supplementary Figure 5). The secondary structure predictions suggested a propensity for both α-helical and β-rich forms (Supplementary Table 3), which aligns the experimental observations in the fibril form.

#### Cyanophlyctin

Cyanophlyctin is secreted on the skin of the amphibian *Euphlyctis cyanophlyctis*, and is potent against both Gram-positive and Gram-negative bacteria, while showing no hemolytic activity around MIC values^59^. Here, the peptide demonstrated relatively potent activity against both *M. luteus* (MIC of 4 μM) and T2-cells (LC_50_ of 18 μM) (Table 1). Structurally, when incubated in D2O over time, cyanophlyctin showed a very subtle secondary structure change, with samples bearing several populations, with a broad peak that included ranges for α, β and disordered structures. In buffer with or without lipids, cyanophlyctin was predominantly in a α-helical fibrillar structure, with a peak in the FTIR spectra range of 1651-1654 cm^−1^ (Figure 4). Correspondingly, the fiber X-ray diffraction pattern resembled cross-α (Figure 3). Specifically, cyanophlyctin displayed reflections at 10 Å and 20 Å spacing, which are orthogonal to reflections at 12 Å and 25 Å spacing. As with PSMα3^8,26,27^, the 10 Å reflection may correspond to the distance between α-helices along the sheet. The orthogonal 12 Å and 25 Å reflections may be attributable to the distance between sheets and pairs of sheets, respectively. The 25 Å reflection can also be attributable to the length of the helix. The secondary structure predictions suggested a propensity for both α-helical and β-rich structures (Supplementary Table 3), which fits the mixed population observed in the fibrillar form incubated in D2O over time. ***Citropin-1.3*.** Citropin-1.3 is secreted from the granular dorsal and submental glands of the Blue Mountains tree-frog *Litoria citropa*^60^. The present analysis showed that it was active against both *M. luteus* (MIC of ∼7 μM) and T2-cells (LC_50_ of ∼35 μM) (Table 1). Structurally, citropin-1.3 displayed a mixed population in the fibril state when incubated in D2O, with a predominant β-rich conformation (Supplementary Figure 5). This aligned with the cross-β polymorph reflected in the fiber diffraction pattern (Supplementary Figure 2). In buffer, citropin-1.3 showed a lipid-induced transition from a major peak at 1622 cm^−1^, indicative of rigid cross-β fibrils, to a major peak at 1651 cm^−1^, indicative of a predominant α-helical structure (Supplementary Figure 5). Citropin-1.3 induced ThT fluorescence (Supplementary Figure 1), which could be attributed to either cross-β or cross-α conformations. The secondary structure predictions suggested a propensity for both α-helical and β-rich forms (Supplementary Table 3), which corroborates the mixed population observed in the fibrillar form.

#### Pleurocidin-like peptide GcSc4C5

Pleurocidin-like peptide GcSc4C5 was identified by analyzing the genomic information and the mRNA transcripts of the *Glyptocephalus cynoglossus* (witch flounder) fish and showed activity against two strains of antibiotic-resistant *Pseudomonas aeruginosa*^61^. Here, the peptide proved a potent antibacterial with an MIC of 2 μM against *M. luteus*, versus relatively moderate activity against T2-cells (LC_50_ of 35 μM) (Table 1). Among the eight ffAMPs that were cytotoxic to T2-cells, pleurocidin-like peptide GcSc4C5 displayed the largest difference between its activity against bacterial and human cells, supporting selectivity for antibacterial activity over cytotoxicity. Pleurocidin-like peptide GcSc4C5 was unstructured in solution, yet the presence of bacterial membrane mimics induced a structural transition toward the α-helical conformation (Supplementary Figure 4). In the fibrillar state, when incubated in D2O, a mixed population formed, with increasing β-rich content over time. The fiber X-ray diffraction pattern taken after several hours of incubation, suggested a cross-β polymorph (Supplementary Figure 3), but also indicated polymorphs with undetermined morphology, which might correspond to the observed mixed population. Specifically, while in one experiment, pleurocidin-like peptide GcSc4C5 incubated for 2.5 h showed a cross-β pattern, in a second experiment, with the same sample incubated for 2 h and 6 h, the diffraction showed different patterns with multiple reflection rings, including at 3.5 Å, 4.3 Å, and 9.2 Å spacings (2 h), or with 4.2 Å, 4.7 Å, 5.1 Å, 6.5 Å and 9.4 Å spacing (6 h), which we could not correlate with a specific known arrangement. When incubated in buffer, pleurocidin-like peptide GcSc4C5 showed predominantly α-helical content (peak at 1651 cm^−1^), and a minor cross-β form (peak at 1622 cm^−1^). Addition of lipids shifted the population towards solely helical fibrils (Supplementary Figure 5). Pleurocidin-like peptide GcSc4C5 induced ThT fluorescence (Supplementary Figure 1), which can be attributed to either cross-β or cross-α conformations. The secondary structure predictions suggested an α-helical propensity (Supplementary Table 3), which failed to align with the experimentally observed β-rich structure in the fibril form.

#### Brevinin-2SKb

Brevinin-2SKb was isolated from the stream brown frog *Rana sakuraii*^62^. The peptide was active against both *M. luteus* (MIC of ∼11 μM) and T2-cells (LC_50_ of ∼40 μM) (Table 1). In solution, it only adopted an α-helical conformation in the presence of bacterial membrane mimics (Supplementary Figure 4). In the fibrillar form, brevinin-2SKb formed a mixed population, without a major transition when incubated in D2O over time, and a predominantly α-helical fibrillar structure content in buffer, with or without lipids (Supplementary Figure 5). Brevinin-2SKb displayed typical cross-β reflection rings in X-ray fiber diffraction (Supplementary Figure 2), which correspond to one of the minor polymorphs observed by the FTIR spectra, yet perhaps the most ordered one with continues structures. The secondary structure predictions suggested a propensity for both α-helical and β-rich structures (Supplementary Table 3), which aligned with the mixed population observed in the fibrillar form.

#### Lasioglossin LL-I

Lasioglossin LL-I was isolated from the venom of the eusocial bee *Lasioglossum laticeps* and shows potent antimicrobial activity against both Gram-positive and Gram-negative bacteria, along with weak hemolytic activity and anticancer activity against PC-12 cancer cells^63^. Our results showed that it was more active against *M. luteus* (MIC of ∼4 μM) as compared to human T2-cells (LC_50_ of ∼45 μM) (Table 1). Lasioglossin LL-I incubated in D2O showed a broad peak in ranges that include α, β and disordered structures, indicating a mixed fibril population with subtle changes over time. In buffer, it was unstructured, with a shift into the random coil/α-helical FTIR spectra range in the presence of lipids (Supplementary Figure 5). The fiber X-ray diffraction pattern indicated that a cross-β fibril exists, and might constitute a minor polymorph (Supplementary Figure 3), and the only one with a repetitive spacing of the peptide stacking, as in the case of brevinin-2SKb. The secondary structure predictions suggested a propensity for both α-helical and β-rich structures (Supplementary Table 3), which fit the mixed population observed in the fibrillar form.

#### Temporin-1Cea

The precursor of temporin-1CEa (Preprotemporin-1CEa) was discovered by molecular cloning of cDNAs of peptide precursors from total RNA extracted from the skin of the Asiatic grass frog *Rana chensinensis*^64^. The mature peptide temporin-1CEa revealed antibacterial activity, along with anticancer activity against MCF7 and Hela cell lines, with 6-20 times more effectiveness compared to hemolysis of human erythrocytes^64^. The present analysis showed that temporin-1CEa had moderate potency against both *M. luteus* (MIC of 31 μM) and T2-cells (LC_50_ of 65 μM) (Table 1). Temporin-1Cea incubated in D2O formed a mixed population in the fibril state, with a time-dependent transition via a α-helical/random coil structure into a solely cross-β structure, with a peak at 1623 cm^−1^ after 72 h (Figure 4). This corresponded with its signature cross-β fiber X-ray diffraction pattern Figure 3). Interestingly, the ThT fibrillation kinetics of temporin-1Cea indicated a decrease in the fluorescence signal after 8-9 h, followed by an increase in fluorescence, up to a steady state (Supplementary Figure 1). This biphasic curve might be indicative of a secondary structure transition from α-helical to β-rich fibrils. When incubated with bacterial and eukaryotic membrane mimics, temporin-1Cea showed a transition from a disordered structure to an α-helical fibril structure with a peak at 1649 cm^−1^ (figure 4). The secondary structure predictions suggested a chameleon propensity (Supplementary Table 3), which fits the mixed population observed in the fibrillar form.

#### Plantaricin-J and Plantaricin-K

Plantaricin-J and plantaricin-K are produced by *Lactobacillus plantarum C11*, a strain of lactic acid bacteria (LAB) present in human microbial flora. They produce class IIb AMPs named two-peptide bacteriocins, which require two different peptides in equal amounts to achieve optimal activity, potentially acting by forming an active complex^65–67^. The two-peptide bacteriocins contain a GxxxG motif known to mediate helix–helix interactions in membrane proteins^66^. Single point mutations replacing glycine in both peptides showed that this motif is critical for their activity^66^, and possibly a mediator of the interaction between the two peptides^66^. In the presence of liposomes, plantaricin-J and plantaricin-K potentially form a complex that assists in structuring the peptides^66^. LAB bacteriocins are known to be very potent, exhibiting activity at pico- to nano-molar concentrations^66^. Our results showed that plantaricin-J and plantaricin-K were active as individual peptides against *M. luteus* with MIC values of ∼8 μM and ∼26 μM, respectively (Table 1). In contrast, plantaricin-J showed weak activity against human T2 cells (LC_50_ of 80 μM) while plantaricin-K was not cytotoxic up to this concentration (Table 1). Correspondingly, LAB bacteriocins are considered safe for human consumption^66^.

Structurally, plantaricin-J showed a predominantly α-helical fibrillar structure in buffer, both in the presence and absence of lipids, while when tested over time in D2O, it showed a mixed population with a broad peak that included ranges of α, β and disordered structures (Supplementary Figure 5). Plantaricin-J displayed a typical cross-β pattern in fiber X-ray diffraction (Supplementary Figure 2), indicating an ordered polymorph with continuous elements. When incubated in D2O over time and in buffer with or without lipids, plantaricin-K formed a mixed population with the major polymorph being of cross-β content (Supplementary Figure 5). This corresponded to the ability of plantaricin-K to induce ThT fluorescence (Supplementary Figure 1), and to its cross-β X-ray fiber diffraction pattern (Supplementary Figure 3). Yet, other polymorphs were also observed. Specifically, in one experiment, the X-ray diffraction of plantaricin-K displayed a cross-β pattern, and in a second experiment, a pattern with reflections at 3.9 Å, 4.3 Å, 5.3 Å and 6.1 Å spacing, which did not correlate with any specific known arrangement (Supplementary Figure 4). The secondary structure predictions suggested an α-helical propensity for plantaricin-K and both α-helical and β-rich forms for plantaricin-J, (Supplementary Table 3), thus failing to predict the experimentally observed β-rich fibril forms of plantaricin-K. It is also possible that the properties of the two peptides will differ when mixed.

#### Cecropin-C

Cecropin-C is produced by *Anopheles gambiae* mosquitoes, which serve as a major malaria vector in Africa^68^. The present analysis showed that cecropin-C had no effect on T2-cells, but potent (MIC of ∼7 μM) against *M. luteus* (Table 1), suggesting high selectivity against bacterial cells. Cecropin-C incubated in D2O showed a mixed fibril secondary structure population, with subtle changes over time. In buffer, it was unstructured, yet shifted towards a random coil/α-helical fibril structure in the presence of lipids (Supplementary Figure 5). The fiber X-ray diffraction pattern indicated that a cross-β fibril exists, which might have comprised a minor polymorph, and the only one with a repetitive spacing of the peptide stacking (Supplementary Figures 2), as in the case of brevinin-2SKb and Lasioglossin LL-I. The secondary structure predictions of cecropin-C suggested both α-helical and β-rich forms in accordance with the experimentally observed mixed population.

#### Aurein-3.3

Aurein-3.3 was isolated from the Southern bell frog *(Litoria/Ranoidea raniformis)* and shows antibacterial and anticancer activity^69^. Our results showed that aurein-3.3 was not active against T2-cells and moderately active (MIC of ∼50 μM) against *M. luteus* (Table 1). When examined over time in D2O, aurein-3.3 showed a random coil/α-helical structure (along with a disordered population) on the first day of incubation, corresponding to its α-helical state in solution (Supplementary Figure 4), and underwent a structural change towards a cross-β structure with a shallow peak at 1621 cm^−1^, after 72 h of incubation (Supplementary Figure 5). The fiber X-ray diffraction pattern of aurein-3.3 taken after several hours of incubation, suggested a cross-β pattern but also indicated other potential fibril morphologies (Supplementary Figure 3). Specifically, aurein-3.3 incubated for 2 h showed a cross-β pattern, and when incubated for one week, showed a pattern with reflections at 4.3 Å, 4.5 Å, 4.7 Å and 5.1 Å spacing, along with other reflections, which we could not have correlated to a specific known arrangement. In buffer, aurein-3.3 was unstructured in its fibrillar form, yet adopted an α-helical structure in the presence of bacterial membrane mimics (Supplementary Figure 5). The secondary structure predictions suggested a propensity for both α-helical and β-rich structures (Supplementary Table 3), which fits the mixed population observed in the fibrillar form.

#### Bombinin H4

Bombinin-H4 is secreted on the skin of the yellow-bellied toad *Bombina variegate*, in two closely related variations, differing only at the first position: one containing isoleucine and the other containing leucine, with the latter used in the current work. Both displayed antibacterial and hemolytic activities^70^. Our results showed that bombinin-H4 was not active against T2-cells and had weak activity (MIC of ∼80 μM) against *M. luteus* (Table 1).

A solution NMR structure of the bombinin-H4 homolog with isoleucine at the first position indicated a helical structure^71^. Previous studies indicated that the addition of dodecylphosphocholine (DPC) micelles to bombinin-H4 in solution induced a shift to α-helical structure, along with a pH-dependent shift of the secondary structure, favoring higher α-helical and β-rich content in acidic and basic pH, respectively^72^. Our resulted showed that bombinin H4 predominantly took on a β-rich structure mixed with some α-helical-rich content in D2O and in buffer, and underwent a structural transition to α-helical content in the presence of eukaryotic membrane mimics, and also in the presence of bacterial membrane mimics yet with a less sharp peak (Figure 4).

The X-ray fiber diffraction of bombinin H4, which exhibited a pattern resembling cross-α with orthogonal reflections at 10 Å and 22 Å spacing (Figure 3), may represent the membrane-active and more stable species of mature fibrils. The 10 Å reflection putatively corresponds to the distance between α-helices along the sheet (axial packing), similar to PSMα3. Another weak reflection at 11 Å spacing might be attributable to the distance between sheets, and the 22 Å spacing, which is orthogonal to 10 Å, might be attributable to the distance between pairs of sheets. Since bombinin H4 contains many glycine residues (25% of the sequence) with no large aromatic residues (Supplementary Tables 2&7), it is possible that the sheets are densely packed with a rather small inter-sheet distance compared to cyanophlyctin. Bombinin H4 is 20-residues in length, with a proline at position 4, thus the 22 Å reflection could correspond to the length of the ordered segment of the amphipathic helix, which is orthogonal to the axial stacking of helices. Bombinin H4 induced ThT fluorescence, which might be attributable to the cross-β as well as the cross-α polymorph, as observed for the cross-α/β-forming uperin 3.5 AMP^8^ and the cross-α-forming PSMα3 cytotoxin^26^. The secondary structure predictions of bombinin H4 suggested an α-helical propensity, corresponding with the cross-α diffraction and the secondary structure in the presence of membranes, but failed to predict the experimentally observed β-rich fibril form, as in the prediction of plantaricin-K.

#### Dolabellanin-B2

Dolabellanin-B2 was isolated from the body-wall, including skin and mucus, of the *Dolabella Auricularia* sea hare and displayed a broad-spectrum antimicrobial activity^73^. Our analysis showed that dolabellanin-B2 was not active against either T2-cells or *M. luteus* up to the tested concentrations (Table 1). Dolabellanin-B2 formed a β-rich structure in solution (Supplementary Figure 4), which was surprising considering its main secondary structure prediction as an amphipathic helix with high hydrophobic vector, yet, indeed, it also displayed a predicted propensity for β structures (Supplementary Table 3). When incubated in D2O over 3 days, dolabellanin-B2 formed a mixed population in the fibrillar state, with helicity appearing after 24 h (Supplementary Figure 5). In buffer, it showed a mixed population, in the presence or absence of lipids, with a predominant cross-β and a minor α-helical content. This corresponded with the cross-β X-ray fiber diffraction pattern (Supplementary Figure 2).

#### Pleurocidin-like peptide WFX

Pleurocidin-like WFX was identified by analyzing the genomic information and the mRNA transcripts of the winter flounder fish *Pseudopleuronectes americanus*; the peptide was also detected on its skin^61,74^. Our results showed that pleurocidin-like peptide WFX was not active against either T2-cells or *M. luteus* up to the tested concentrations (Table 1). Despite its definition in the public databases as an antimicrobial (Supplementary Table 4), no reports of such antimicrobial activity were found^61,74^, which might indicate that this peptide has a different function on the skin of the fish. Structurally, pleurocidin-like peptide WFX formed a mixed population in the fibrillar state, with a major β-rich content when incubated in D2O (Figure 4), which corresponded with its cross-β X-ray fiber diffraction pattern (Figure 3). When incubated in buffer, with or without lipids, pleurocidin-like peptide WFX showed a predominant β-rich structure and a minor α-helical content (Figure 4). The secondary structure predictions suggested a propensity for both α-helical and β-rich structures (Supplementary Table 3), which aligns with the mixed population observed in the fibrillar form.

### Amino acid prevalence in the ffAMPs

To complement the structural analysis of the 14 ffAMPs, a sequence analysis of their amino acid prevalence was conducted, and compared to the prevalence in short (<40 residues) AMPs in the entire CAMP_R3_ database. The analysis showed that lysine and glycine residues were most prevalent among the 20 amino acids in the 14 ffAMPs, with proline and threonine proving the least prevalent (Supplementary Table 7). Proline is likely rare due to its critically disruptive effect on secondary structures. It was also much less prevalent in the 14 ffAMPs compared to short AMPs in the entire database, suggesting a selected-against property. Threonine and cysteine residues also showed lower prevalence in the 14 ffAMPs compared to short AMPs in the entire database. Interestingly, single and an odd number of cysteines were previously shown to be rare in short helical AMPs compared to AMPs in general, putatively due to their effect on supramolecular assemblies via inter-molecular disulfide bonds^39^.

Grouping amino-acids by physico-chemical properties (Supplementary Table 8) showed that the 14 ffAMPs were especially rich in non-polar residues compared to polar residues, and positively charged residues were much more abundant compared to negatively charged residues. This aligns with the mechanism of action of ffAMPs, which required interactions with the negatively charged bacterial membranes. Correspondingly, structures of α-helical, toxic, fibril-forming peptides displayed hydrophobic interfaces between amphipathic helices^8,26,27,40^ and hydrophobic patches, with protruding positively charged amino acids, on the fibril surface^39,40^.

Of note, among the ffAMPs, pleurocidin-like peptide WFX, dolabellanin-B2, and bombinin H4 contained the lowest percentage of lysine and arginine residues (Supplementary Table 7). Pleurocidin-like peptide WFX also showed the highest percentage of glutamate and aspartate residues, with an overall side chains net charge of zero. Considering AMPs interactions with negatively charged membrane lipids, the prevalence of charged residues directly corresponds to the activity level against *M. luteus*, with pleurocidin-like peptide WFX, dolabellanin-B2, and bombinin H4 being the least toxic (Table 1).

## DISCUSSION

This work applied a bioinformatics platform to identify antimicrobial peptides that form fibrils with amyloid properties, based on predictions of secondary structure and hydrophobic moment. Short toxic peptides with amyloid-forming capacities were widespread across organisms. Fourteen of the 26 AMPs tested showed fibril formation as well as amyloid properties reflected in fiber X-ray diffraction spectra, with five also demonstrating ThT amyloid dye binding. Of note, not all amyloids bind ThT, which might be related to alteration in the cavities along the fibril^26^. Four ffAMPs (cupiennin-1, pleurocidin-like peptide GcSc4C5, temporin-1Cea, and aurein-3.3) showed a time-dependent transition to β-rich fibrils, which is typical for amyloids, two of which (pleurocidin-like peptide GcSc4C5 and temporin-1Cea) also bound ThT (Table 1).

### Lipid-induced structural transition of ffAMPs in solution and/or in the fibril form

It is well known that AMPs can undergo conformational changes towards α-helical structures upon contact with membrane lipids^75–78^. The present work demonstrated that this secondary structure chameleon switch can occur in solution and in the fibril state. Five ffAMPs showed lipid-induced structural transition toward an α-helical conformation in solution (Supplementary Table 5 and Supplementary Figure 4). Four of those, i.e., citropin-1.3, bombinin H4, temporin-1Cea and pleurocidin-like peptide GcSc4C5, showed a lipid-induced shift to a more α-helical content in the fibrillar state as well (Figure 4 and Supplementary Figure 5). Interestingly, these four ffAMPs also induced ThT fluorescence. Three additional ffAMPs, aurein 3.3, cecropin-C, and lasioglossin LL-I, showed lipid-induced structure transition in the fibrillar form only, and did not induce ThT fluorescence. Taken together, a correlation may exist between ThT amyloid dye binding and the chameleon property of lipid-induced secondary structure transition both in solution and the fibrillar form. This aligns with our previous report of ThT binding and lipid-induced transition of uperin 3.5 AMP^8^. The high-resolution structures of both the cross-α and cross-β fibrils of uperin 3.5^8^ ^32^ validated and visualized the chameleon secondary structure switch properties of amyloid fibrils; here we revealed the extent of this aptitude in natural sequences from a variety of organisms.

### Time-dependent structural transition in the fibril form

While the lipid-induced structure switch resulted in higher α-helical content, time-dependent secondary structure transition indicated polymorphism within the sample in the same conditions (Figure 4 and Supplementary Figure 5). Cupiennin-1, temporin-1Cea, pleurocidin-like peptide GcSc4C5 and aurein-3.3 showed a transition from α-helical or a mixed population towards a predominantly β-rich population, thereby resembling cross-β amyloids. Dolabellanin-B2 mostly remained in the β-rich form, yet a α-helical population arose over time. This β to α transition is rare among tested AMPs. A change in the distribution of the citropin-1.3, bombinin H4 and pleurocidin-like WFX populations was observed over time, yet without a specific trend towards a particular secondary structure form. Notably, although the main FTIR-measured polymorph of bombinin H4 was the β-form, X-ray fibril diffraction indicated a pattern that resembled the cross-α, with sharp crossed arcs. Bombinin H4 also induced ThT binding, which might be attributed to the cross-α form, similar to PSMα3^26,27^, or to the mixed cross-α/β forms as demonstrated for uperin 3.5^8^. Of note, exploring the secondary structure of different PSMs using synchrotron radiation circular dichroism spectroscopy and FTIR revealed a mixture of α and β structures within the same sample^79^ similar to our observations for the ffAMPs. Specifically, for PSMα3, although the major population was α-helical, a minor population with β-rich content was observed. This observation was recently validated by two-dimensional infrared (2DIR) studies, which identified a peak related to cross-β structures, appearing after four days of incubation, in addition to the main peak attributed to α-helical content^55^. This suggests that the chameleon secondary structure switch is a widespread phenomenon, inherent to many self-assembling polymorphic sequences forming toxic amyloid fibrils. This property might be related to obtaining control over stability, adherence, activity levels, and specificity.

### α-helical fibril structure correlates with cytotoxicity

As shown in Table 1, all eight ffAMPs that were toxic against human T2-cells were α-helical in the fibril form or adopted α-helical conformation in the presence of lipids. Of the six ffAMPs which were not active against T2-cells, cecropin-C showed potent activity against *M. luteus* and transitioned into the α-helical fibril form in the presence of lipids. The other five ffAMPs, displayed different secondary structures. Specifically, plantaricin-K, dolabellanin-B2 and pleurocidin-like WFX showed mixed α/β fibril forms, which were insensitive to the presence of lipids. Bombinin H4 showed a transition from a predominantly β-rich form to α-helical fibril form in the presence of lipids, especially those mimicking bacterial cells. It is possible that the β-rich fibrils which are still present hinder activity, leading to the high MIC (∼80 μM) against *M. luteus*. Aurein-3.3 was unstructured in the fibril form and transitioned to α-helical fibrils in the presence of bacterial members, suggesting an α-helical antibacterial form, with a moderate MIC of ∼50 μM. Overall, these findings support the hypothesis that α-helicity in the fibril form is more relevant to activity^8,26,27^. It is possible that time-dependent transition to a β-rich fibril, as observed for cupiennin-1, pleurocidin-like peptide GcSc4C5, temporin-1Cea and aurein-3.3, serves as a mechanism for storage of peptides awaiting the encounter with target cells.

One of the major challenges of amyloid systems is understanding their fibrillation-dependent toxicity mechanisms. Namely, studying the dynamics between monomers, oligomers and fibrils, and their role in amyloid toxicity. This is evidenced by the lack of clarity regarding mechanisms of neurotoxicity in human neurodegenerative diseases, despite dozens of research years. This perplexity is further complicated by the extensive polymorphs in the fibrillar state, to the extreme of fibrils showing populations with different secondary structures. We previously hypothesized that PSMα3, a cross-α toxic amyloid peptide, co-aggregates with cell membranes in a process requiring the constant presence of monomers and the ability to form the fibrils, suggesting that toxicity can be viewed as a complex process rather than attributed to a specific toxic entity^8,26,27^. Here we substantiate time- and lipid-induced secondary structure change in the fibril form of ffAMPs, which further complicates the analysis of population dynamics in relation to toxicity mechanisms. Overall, we conjecture that both cross-α/β configurations are subject to a dynamic assembly-disassembly equilibrium that underlies their toxic properties, putatively via dynamic co-aggregation with the membrane.

### High-resolution structure of aurein-3.3 reveals cross-β with kinked β-sheets

High-resolution structure determination of aurein-3.3 using cryogenic electron microscopy (cryo-EM)^32^ revealed a cross-β fibril with an arrangement comprised of six peptides per fibril layer, all showing kinked β-sheets, yielding a compact, rounded, fibril. Such sharp kinks in the b-strand backbone were suggested as the hallmark of low-complexity amyloid-like reversible kinked segments (LARKS) within functional amyloids^80–86^. The structure reinforced the secondary structure predictions (Supplementary Table 3), and the experiments (Table 1), including the X-ray fibril diffraction spectra showing cross-b fibrils, and the FTIR measurements showing a time-dependent shift to the b-form. The mixed populations of aurein-3.3, however, might be related to the liable properties of fibrils composed of kinked b-sheets. Recently, it was proposed that both kinked and extended β-strands contribute to fibril instability, which might be an embedded feature to allow amyloid reversibility ^87^. Labile and reversible amyloid fibril formation was suggested to enable a wide range of functionality in various physiological contexts^88^. Indeed, most ffAMPs studied here displayed substantial polymorphism in the secondary structure of the fibril state, within the same sample, and with time- or lipid-induced shift of population dynamics, suggesting it as an essential property of functional amyloids, specifically of those involved in toxic activities.

### The amyloid-antimicrobial link

The prevalence of ffAMPs displaying amyloid properties aligns with previous reports of other amyloid-like AMPs^3–10^ and complements the observation that human amyloids associated with neurodegenerative and systemic diseases possess antimicrobial properties^11,13–16,20,21,23–25,33^. This amyloid-antimicrobial link assigns a physiological role in neuro-immunity to amyloids, thought, to date, to be exclusively pathological ^33^. Here we demonstrate the extensivity of the amyloid property in AMPs and our ability to identify amyloid-forming AMPs based on secondary structure predictions and physiochemical properties. This overall provides substantial support for the amyloid-antimicrobial link and hypotheses regarding the roles of pathological amyloids in host defense.

### Amyloid fibrillation of AMPs as a mechanism to enhance stability and adherence

Nine out of the 14 ffAMPs examined here (64%) are produced by marine or amphibian organisms, which amounted to a 1.7-fold enrichment compared to the prevalence of short AMPs (<100 residues) in the entire CAMPR3 database produced by marine or amphibian organisms^36^ (Supplementary Table 9). This suggests advantageous value of amyloid fibril formation by AMPs in organisms living in marine ecosystems and other distinctive environments. The known mechanical and chemical stability, and adherence properties of amyloids^89^ can open paths to develop antimicrobials with enhanced oral bioavailability, stability under harsh conditions, adherence to surfaces, and longer shelf-life.

## CONCLUSIONS

Here we demonstrate the identification of 14 amyloid-forming AMPs via a sequence-based bioinformatics platform. The presented findings reinforce the structural and functional connection between antimicrobial activity and amyloidogenicity. A prominent feature of most of these AMPs is their mixed secondary structure population in the fibrils form, including time- and lipid-dependent transitions. Cytotoxic activities were correlated with a fibrillar helical secondary structure. Ordered self-assembly could have been evolved in AMPs across kingdoms of life for functional utilization of amyloid stability and adherence properties, along with regulation of storage versus activity roles, via fibril secondary structure switch.

## ASSOCIATED CONTENT

### Supporting Information

Sequence-based bioinformatic parameters, thresholds, and numerical results of AMPs displaying amphipathic helical and chameleon properties; Sequences and reporting citations of the 14 ffAMPs; Physio-chemical properties and secondary structure prediction of the 14 ffAMPs; Source organisms and identification codes of the 14 ffAMPs; Database identification codes of the 14 ffAMPs; Secondary structure analyses of the ffAMPs in solution using circular dichroism (CD); Amino acid prevalence in the 14 ffAMPs; Amino acid prevalence classified by physicochemical properties; Prevalence of AMPs produced by marine organisms and amphibians; Fibrillation kinetics of five ffAMPs that bound thioflavin T (ThT); X-ray fiber diffraction of cross-β forming ffAMPs; X-ray fiber diffraction ffAMPs displaying structural changes; Secondary structure analyses of the ffAMPs in solution, as measured by circular dichroism; Secondary structure FTIR analyses of the ffAMPs in the solid/ fibrillar form (PDF).

## Notes

The authors declare no competing financial interest.

## ACKNOWLEDGMENT

The synchrotron X-ray fiber diffraction data collection experiments were performed at beamline P14, operated by EMBL Hamburg at the PETRA III storage ring (DESY, Hamburg, Germany). We are grateful to Gleb Bourenkov and the teams at EMBL Hamburg for their assistance. We acknowledge guidance and support from Yaron Kauffmann from the MIKA Electron Microscopy Center of the Department of Materials Science & Engineering at the Technion and from Na’ama Koifman from the Russell Berrie Electron Microscopy Center of Soft Matter at the Technion, Israel. We acknowledge support from the Ilse Katz Institute for Nanoscale Science and Technology, Ben Gurion University of the Negev, Israel.

## ABBREVIATIONS

AMPs: antimicrobial peptids
ffAMPs: fibril-forming antimicrobial peptides
Aβ: amyloid-β
PSMα3: phenol-soluble modulin α3
CAMP_R3_: collection of anti-microbial peptides
aa: amino acids
µH: hydrophobic moment
DMSO: dimethyl sulfoxide
HFIP: 1,1,1,3,3,3-hexafluoro-2-propanol
ThT: thioflavin T
D2O: deuterium oxide
LB: Luria-Bertani
DOPE: 1,2-dioleoyl-sn-glycero3-phosphoethanolamine
DOPG: 1,2-dioleoyl-sn-glycero-3-phospho- (1’-racglycerol)
DOPC: 1,2-dioleoyl-sn-glycero-3-phosphocholine
Chol: cholesterol
SM: sphingomyelin
UPddw: Ultra-pure double distilled water
MIC: Minimal inhibitory concentration
*M. luteus*: *Micrococcus luteus*
TEM: Transmission electron microscopy
MIC: minimal inhibitory concentrations
SUV: Small unilamellar vesicle
LDH: lactate dehydrogenase
LC50: 50% lethal concentration
CD: Solution circular dichroism
ATR: attenuated total internal reflections
FTIR: fourier transform infrared
HCl: hydrochloric acid
TFA: trifluoroacetic acid
3D: three-dimensional

## SUPPLEMENTARY MATERIALS

**Supplementary Table 1.** Sequence-based bioinformatic parameters, thresholds, and numerical results of AMPs displaying amphipathic helical and chameleon properties.

|  |  | AMPs with unknown 3D structures |  |  | AMPs with known 3D structures |  |  |
| --- | --- | --- | --- | --- | --- | --- | --- |
|  |  | Iteration A | Iteration B | Iteration C | Iteration A | Iteration B | Iteration C |
| Number of AMPs in database |  | 2916 |  |  | 105 |  |  |
| Minimal prediction length |  | 6aa | 5aa | 5aa | 6aa | 5aa | 5aa |
| Maximal gap length |  | 3aa |  |  |  |  |  |
| Minimal α-helix score Jpred | | 7.5 | 7 | 7 | 7.5 | 7 | 7 |
| Maximal difference between helix and beta score Tango |  | - | 3 | 3 | - | 3 | 3 |
| Minimal μHscore Jpred | | 0.8 | 0.8 | 0.7 | 0.8 | 0.8 | 0.7 |
| Minimal μHscore Tango | | 0.8 | 0.61 | 0.61 | 0.8 | 0.61 | 0.61 |
| AMPs predicted as α-helical | Number of AMPs | 1690 | 1837 | 1837 | 61 | 65 | 65 |
| | Number of AMPs after applying the μH score threshold | 26 | 44 | 122 | 4 | 4 | 23 |
| AMPs predicted as chameleons | Number of AMPs | 2287 | 1447 | 1447 | 47 | 20 | 20 |
| | Number of AMPs after applying the μH score threshold | 19 | 181 | 181 | 8 | 9 | 9 |

The number of AMPs shorter than 100 amino acids from the CAMPR3 database passing the indicated threshold values set in three different iterations are described for two groups: AMPs with a known three-dimensional (3D) structures and AMPs with unknown 3D structures. μH denotes for hydrophobic moment. "-" denotes that threshold values were not set. The Jpred server was used to determine α-helical reliability scores and the Tango server was used to determine reliability scores of α-helical and $\beta$ -structure predictions of chameleon AMPs. In iteration A, threshold values were set according to "sensible" criteria. Next, secondary structure predictions were compared to experimental results in the group of AMPs with known 3D structures, and the μH value of chameleon predictions was compared to that of uperin 3.5 to optimize thresholds for iteration B. In iteration C, threshold values were further optimized based on experimental validation of fibril formation.

**Supplementary Table 2.**
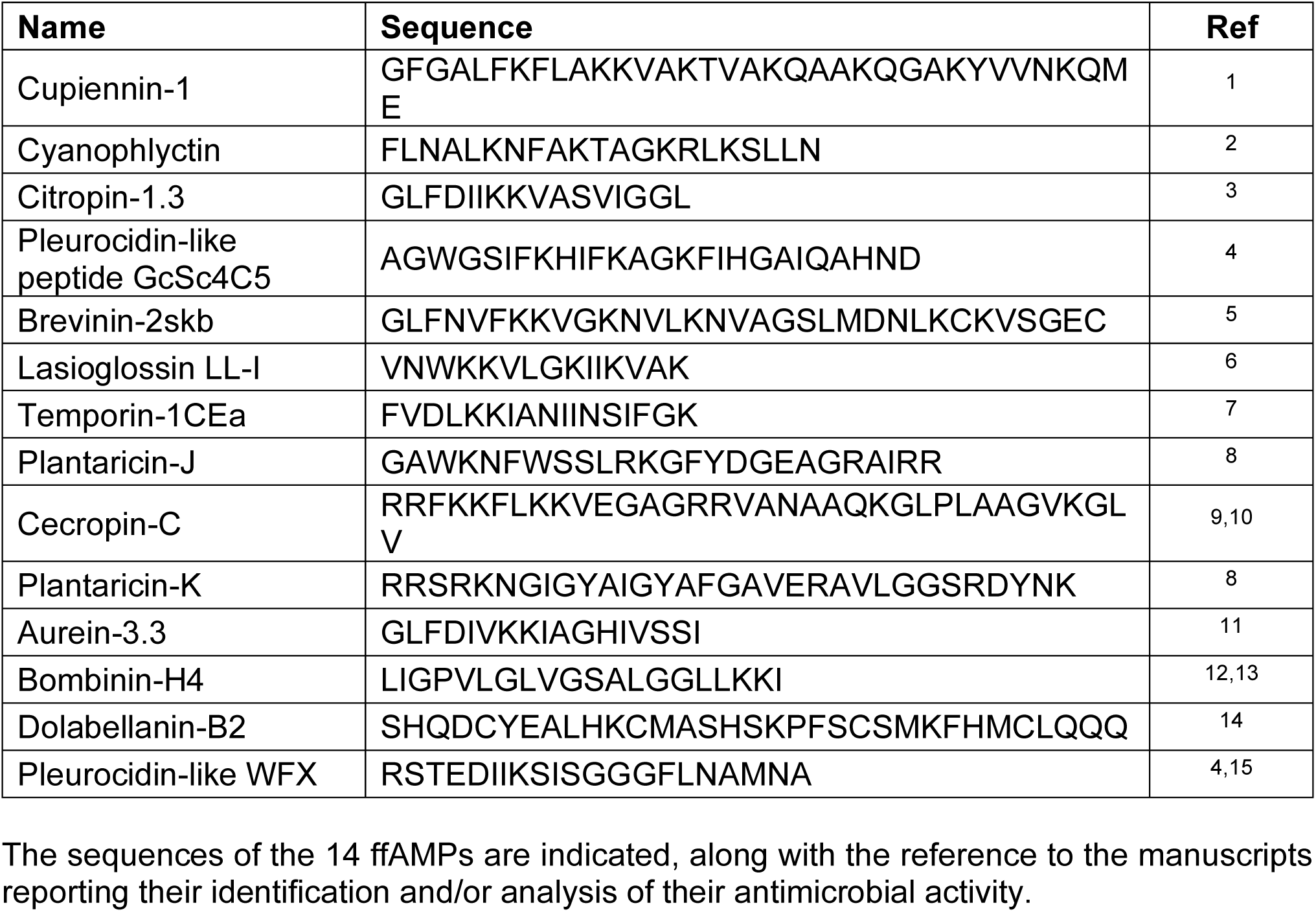
Sequences and reporting citations of the 14 ffAMPs.

**Supplementary Table 3.**
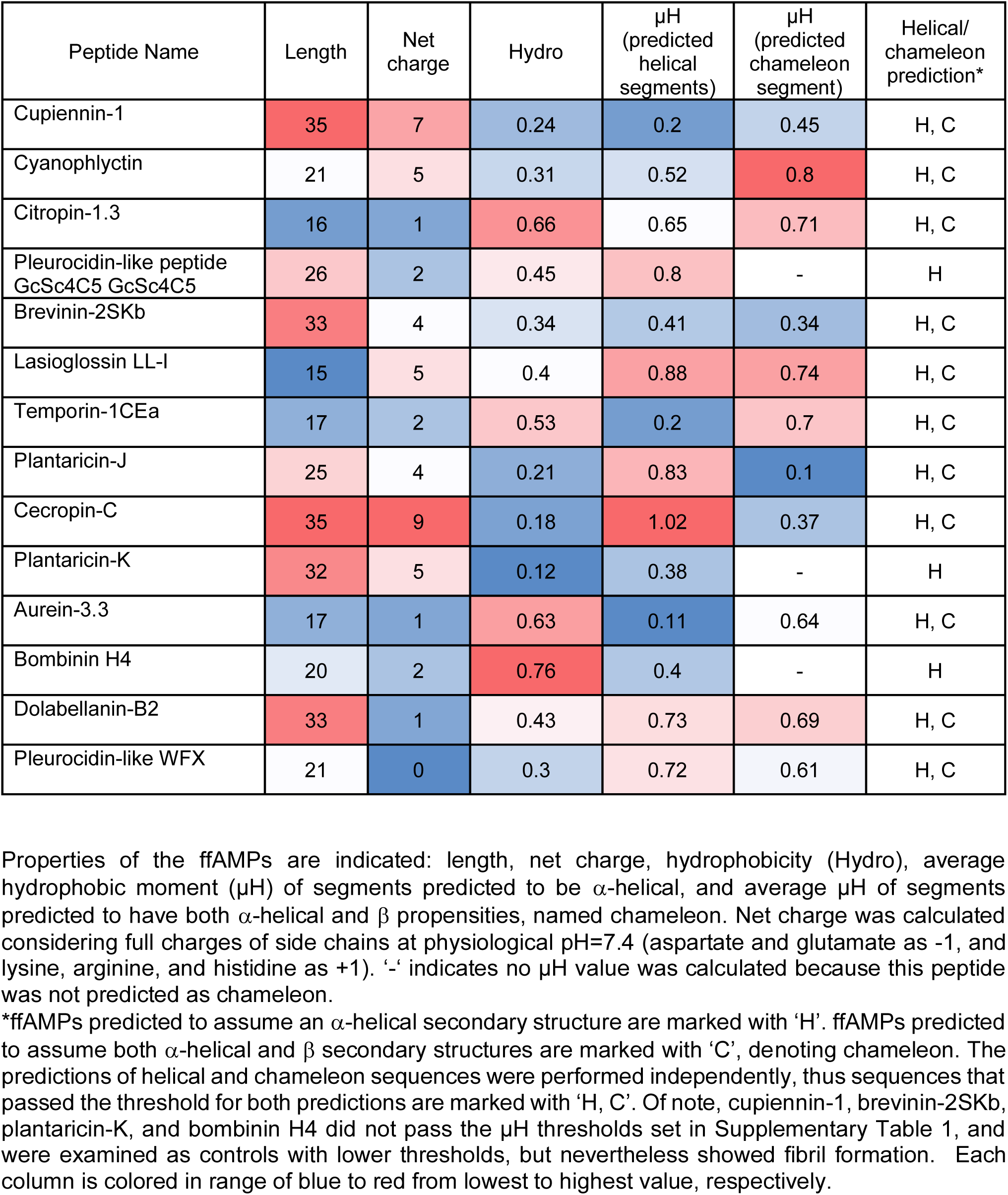
Physio-chemical properties and secondary structure prediction of the 14 ffAMPs.

**Supplementary Table 4.**
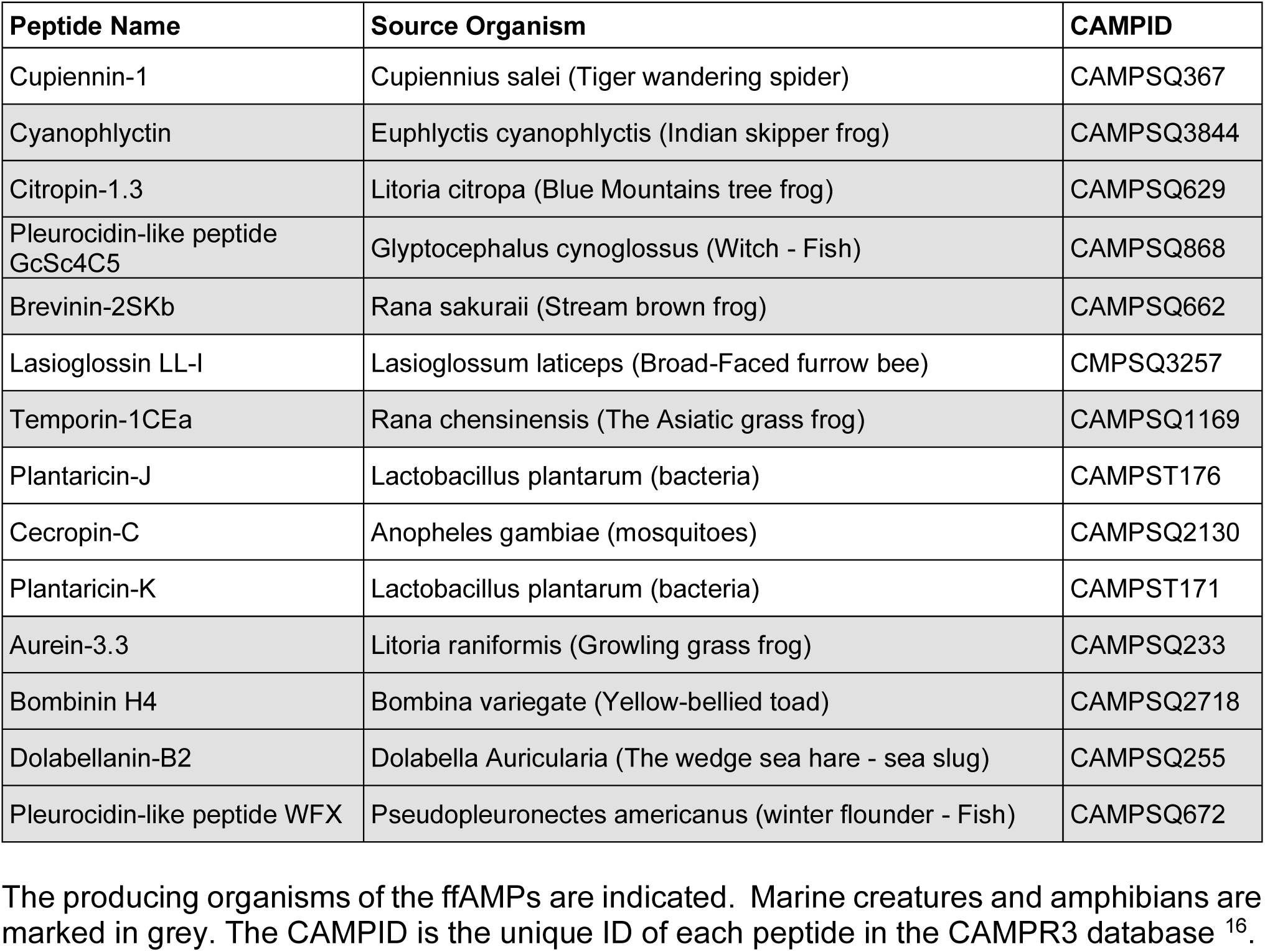
Source organisms and identification codes of the 14 ffAMPs.

**Supplementary Table 5.**
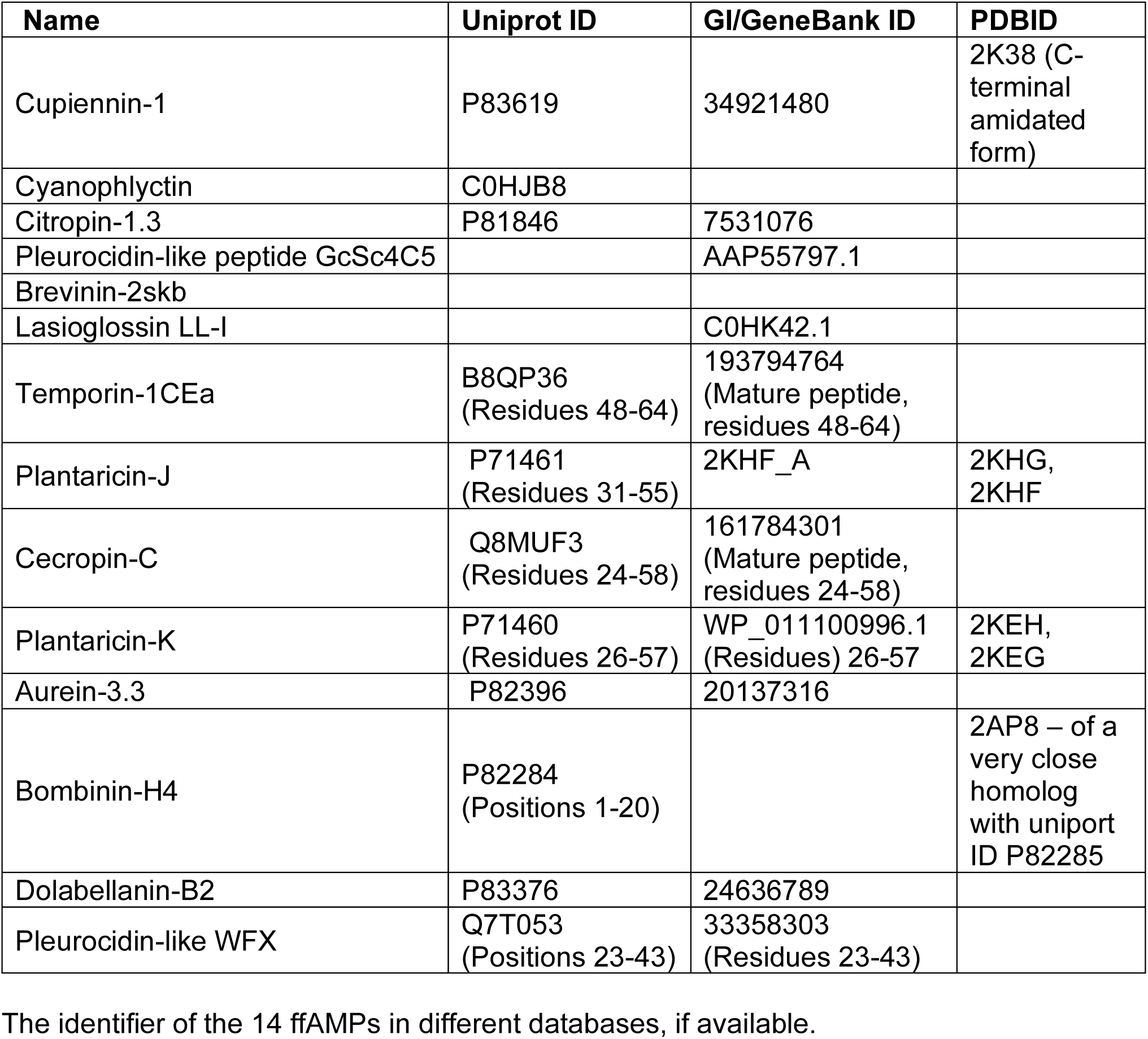
Database identification codes of the 14 ffAMPs.

**Supplementary Table 6.**
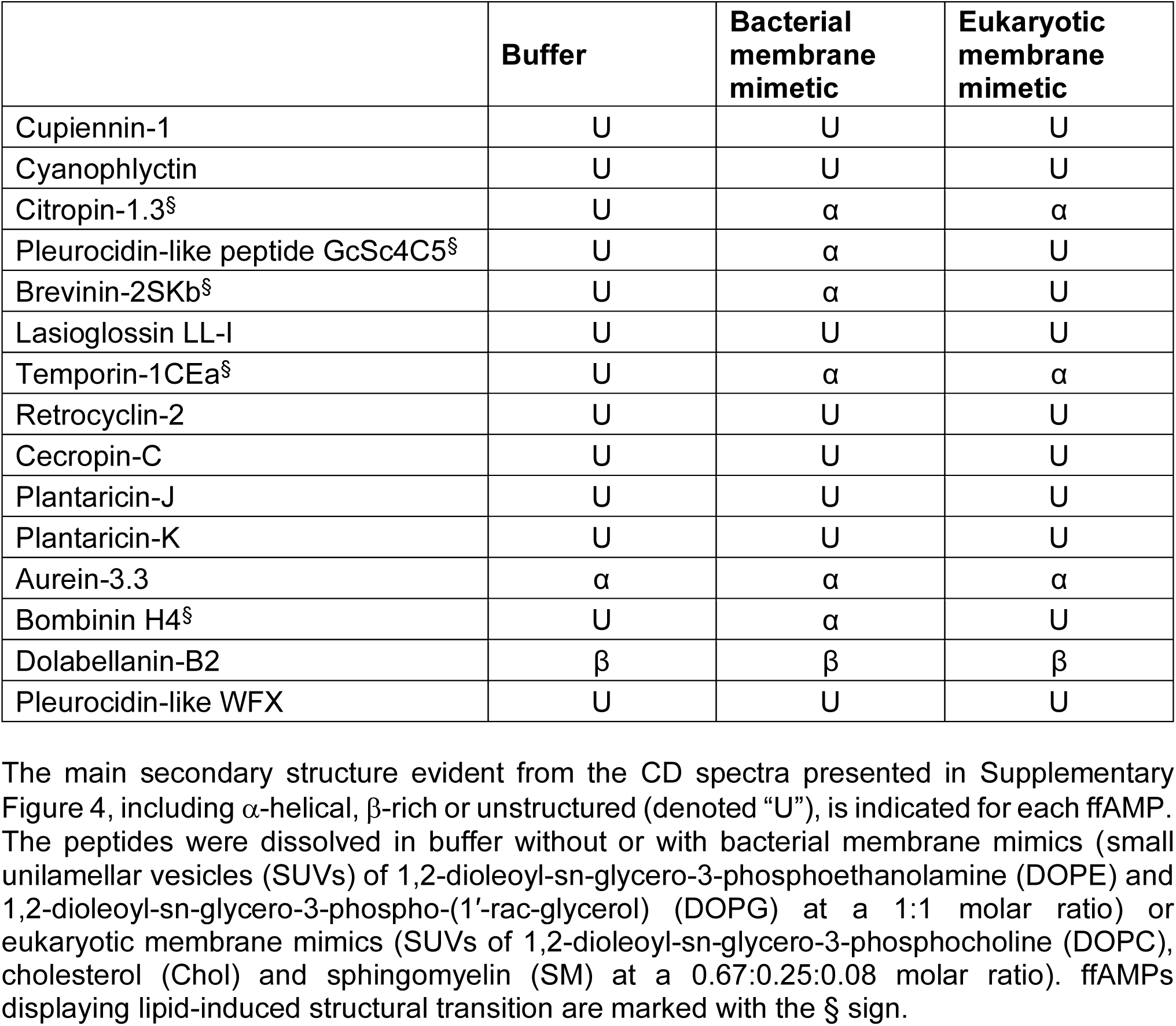
Secondary structure analyses of the ffAMPs in solution using circular dichroism (CD)

**Supplementary Table 7.**
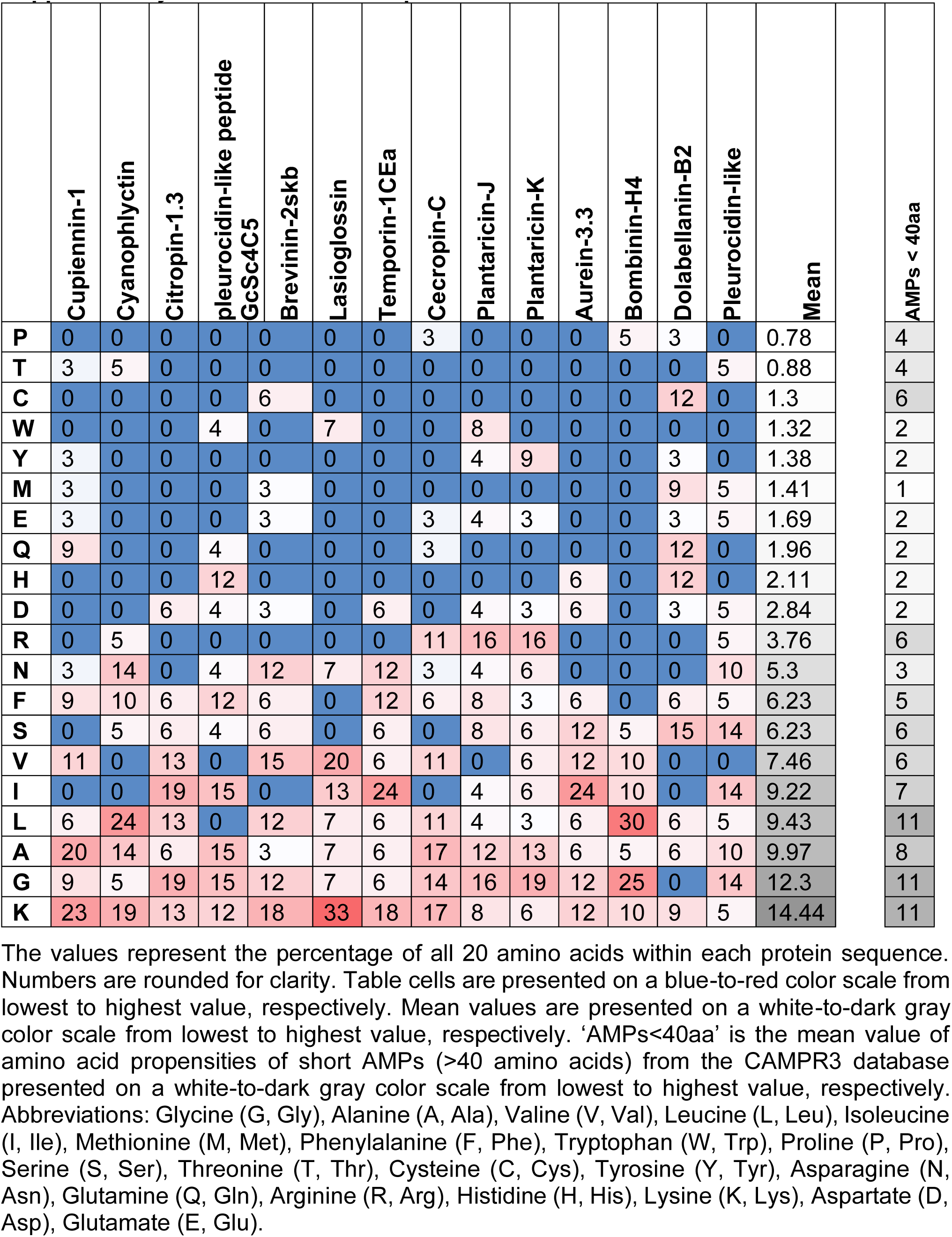
Amino acid prevalence in the 14 ffAMPs.

**Supplementary Table 8.**
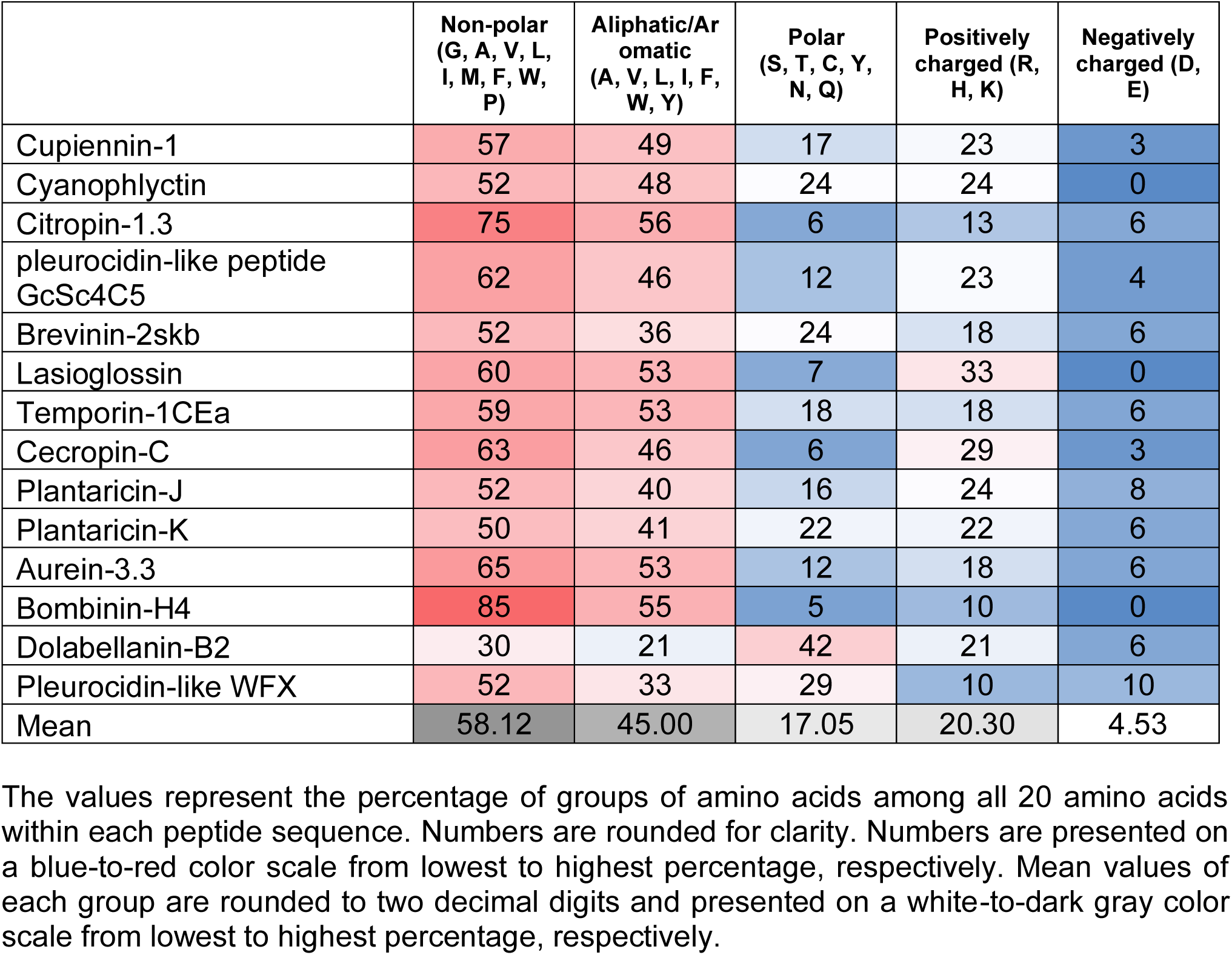
Amino acid prevalence classified by physicochemical properties.

**Supplementary Table 9.**
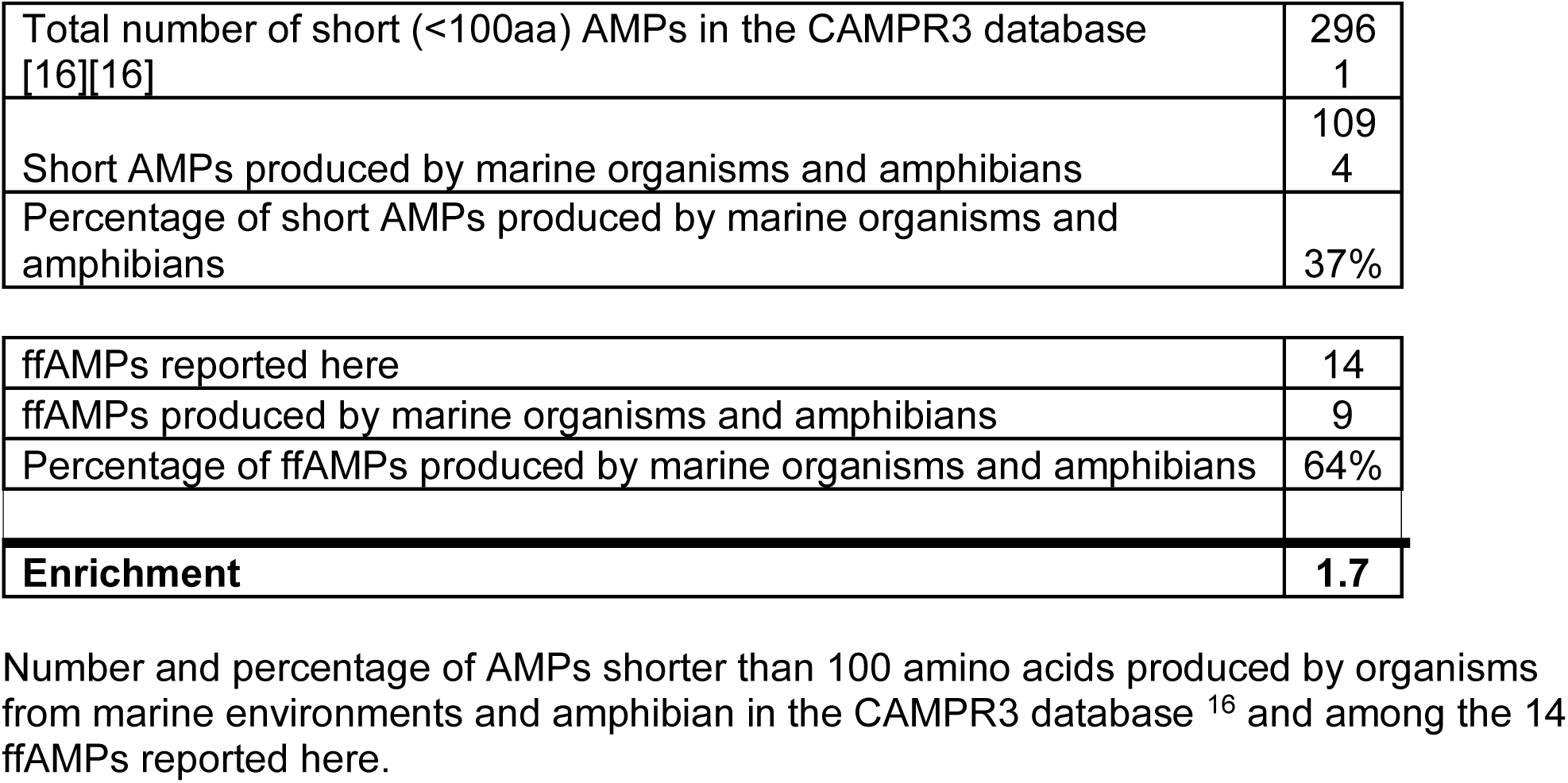
Prevalence of AMPs produced by marine organisms and amphibians.

|  |  |
| --- | --- |
| Total number of short (<100aa) AMPs in the CAMPR3 database<br>[16][16] | 296<br>1 |
| Short AMPs produced by marine organisms and amphibians | 109<br>4 |
| Percentage of short AMPs produced by marine organisms and amphibians | 37% |
| ffAMPs reported here | 14 |
| ffAMPs produced by marine organisms and amphibians | 9 |
| Percentage of ffAMPs produced by marine organisms and amphibians | 64% |
| **Enrichment** | **1.7** |

Number and percentage of AMPs shorter than 100 amino acids produced by organisms from marine environments and amphibian in the CAMPR3 database <sup>16</sup> and among the 14 ffAMPs reported here.

**Supplementary Figure 1.**
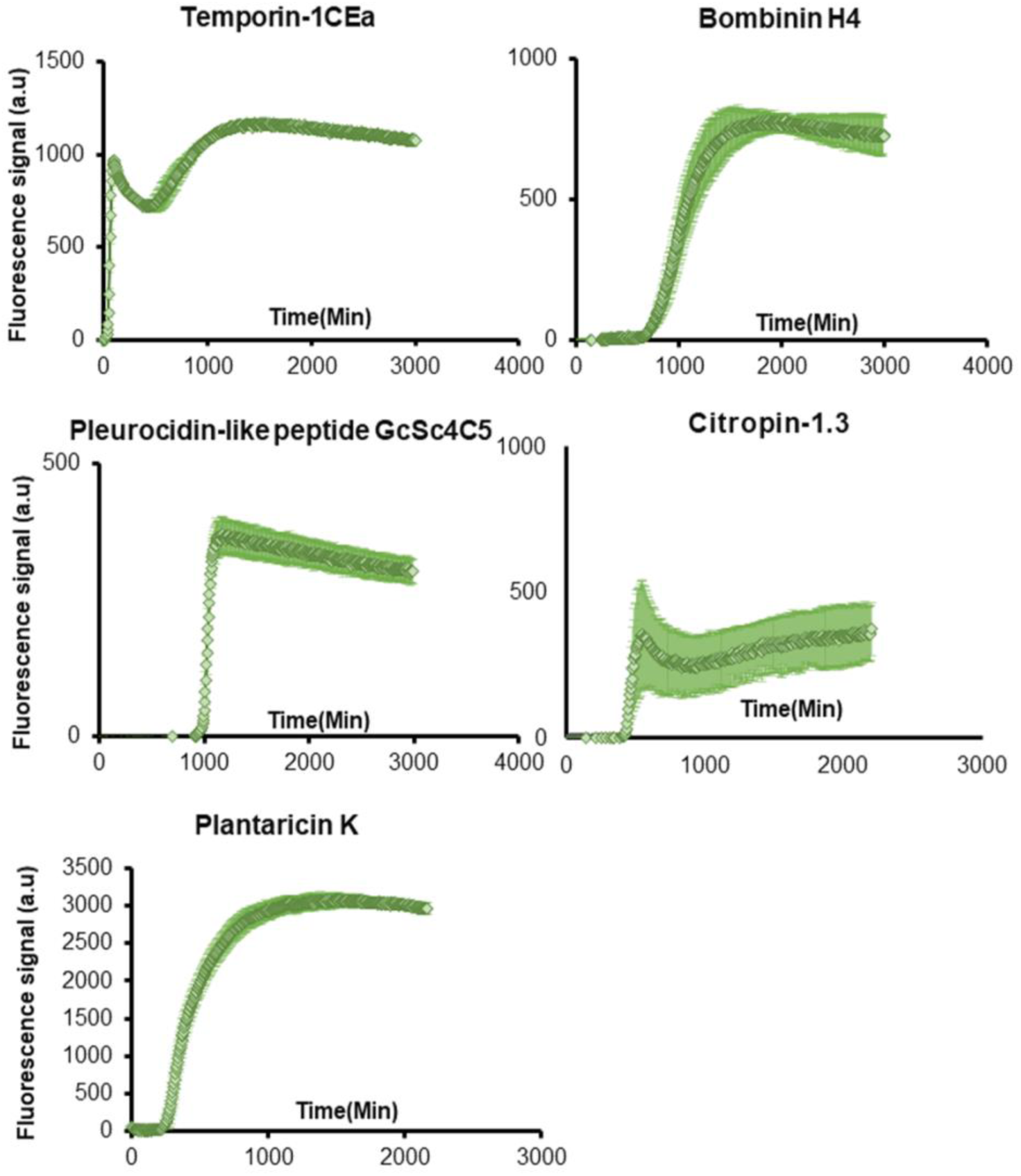
Fibrillation kinetics of five ffAMPs that bound thioflavin T (ThT) Fibrillation kinetics of 100 µM temporin-1CEa, bombinin H4, pleurocidin-like peptide GcSc4C5, citropin-1.3, and plantaricin-K monitored by thioflavin-T (ThT) binding. The graphs show the mean fluorescence readings of triplicate ThT fluorescence measurements over time. Error bars represent standard errors of the means.

**Supplementary Figure 2.**
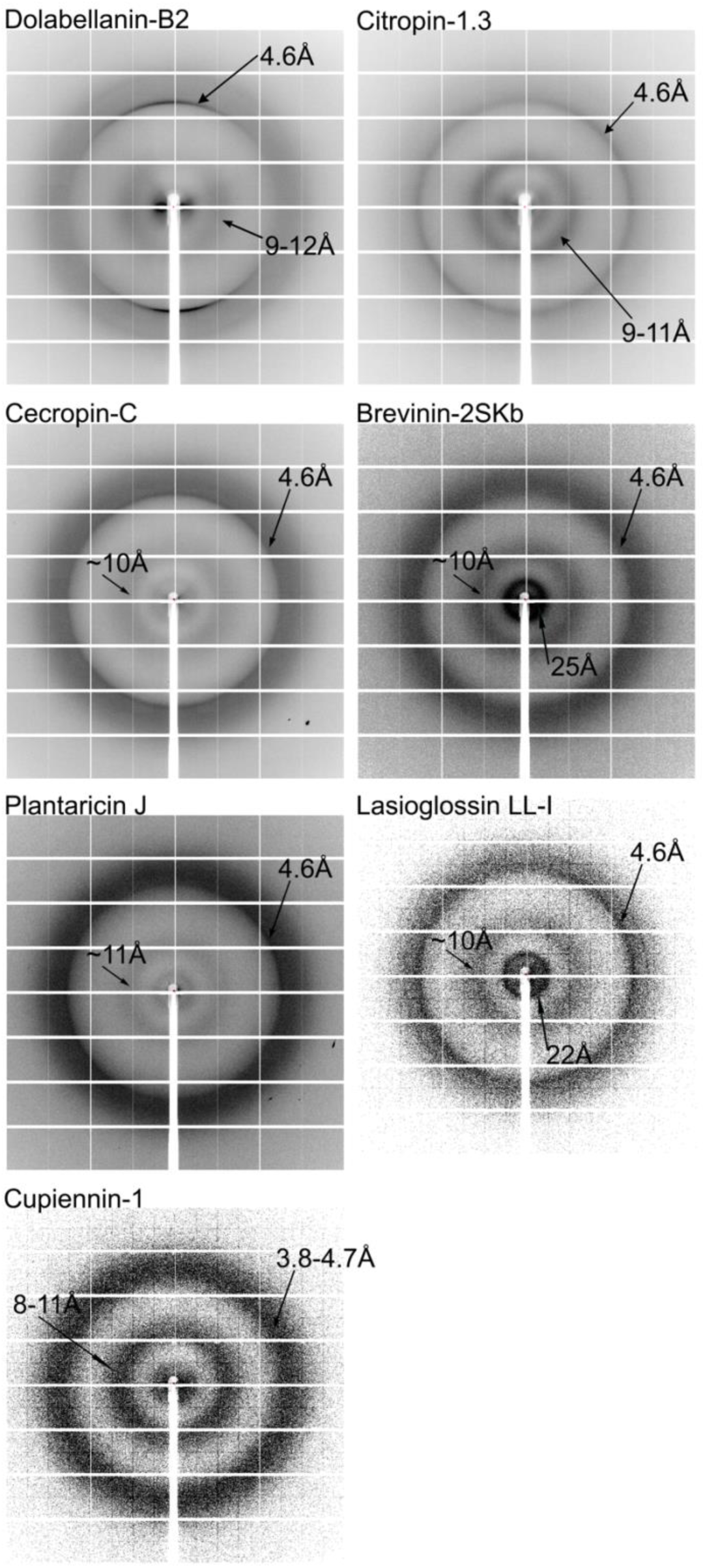
X-ray fiber diffraction of cross-β forming ffAMPs. The X-ray fiber diffraction of ffAMPs showed cross-β diffraction patterns. Major reflections are indicated. All ffAMPs were incubated in double distilled water at 10 mM. Dolabellanin-B2 was incubated for 2 h. Citropin-1.3 was incubated for 2 h. Cecropin-C was incubated for 2.5 h. Brevinin-2SKb was incubated for 2.5 h. Plantaricin-J was incubated for 2.5 h. Lasioglossin LL-I was incubated for 2.5 h. Cupiennin-1 was incubated for 2.5 h, and showed very diffused rings, including around 4.7 Å, which is usually sharp in cross-β amyloids, and was thus not classified as cross-β in Table 1.

**Supplementary Figure 3.**
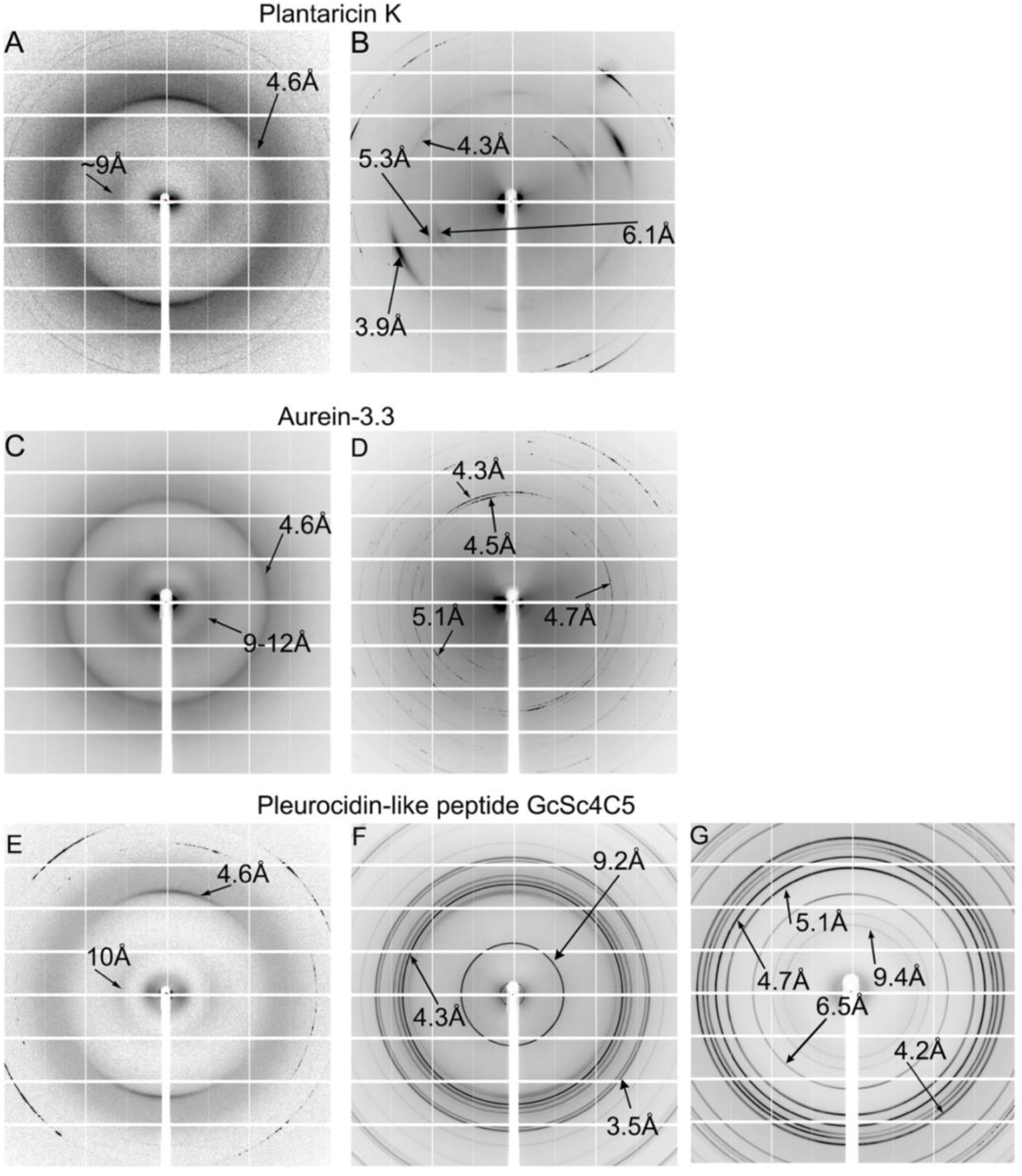
X-ray fiber diffraction ffAMPs displaying structural changes. The X-ray fiber diffraction patterns of ffAMPs showed time-dependent or seemingly random structural changes. Major reflections are indicated. All ffAMPs were incubated in double distilled water at 10 mM. **A&B**. Plantaricin-K was incubated for 2.5 h (A) and 2 h (B). Peptide incubation and X-ray measurements were performed on separate days. **C&D**. Aurein 3.3 was incubated for 2 h (C) and for 7 days (D), with X-ray measurements performed on the same day. **E-G**. Pleurocidin-like peptide GcSc4C5 was incubated for 2.5 h (E), 2 h (F) and 6 h (G). The setup and X-ray measurements of samples shown in panels F&G were performed on the same day. Measurements of samples shown in panel E were performed on a separate day. The diffraction in panels A, C and E indicate a cross-β pattern.

**Supplementary Figure 4.**
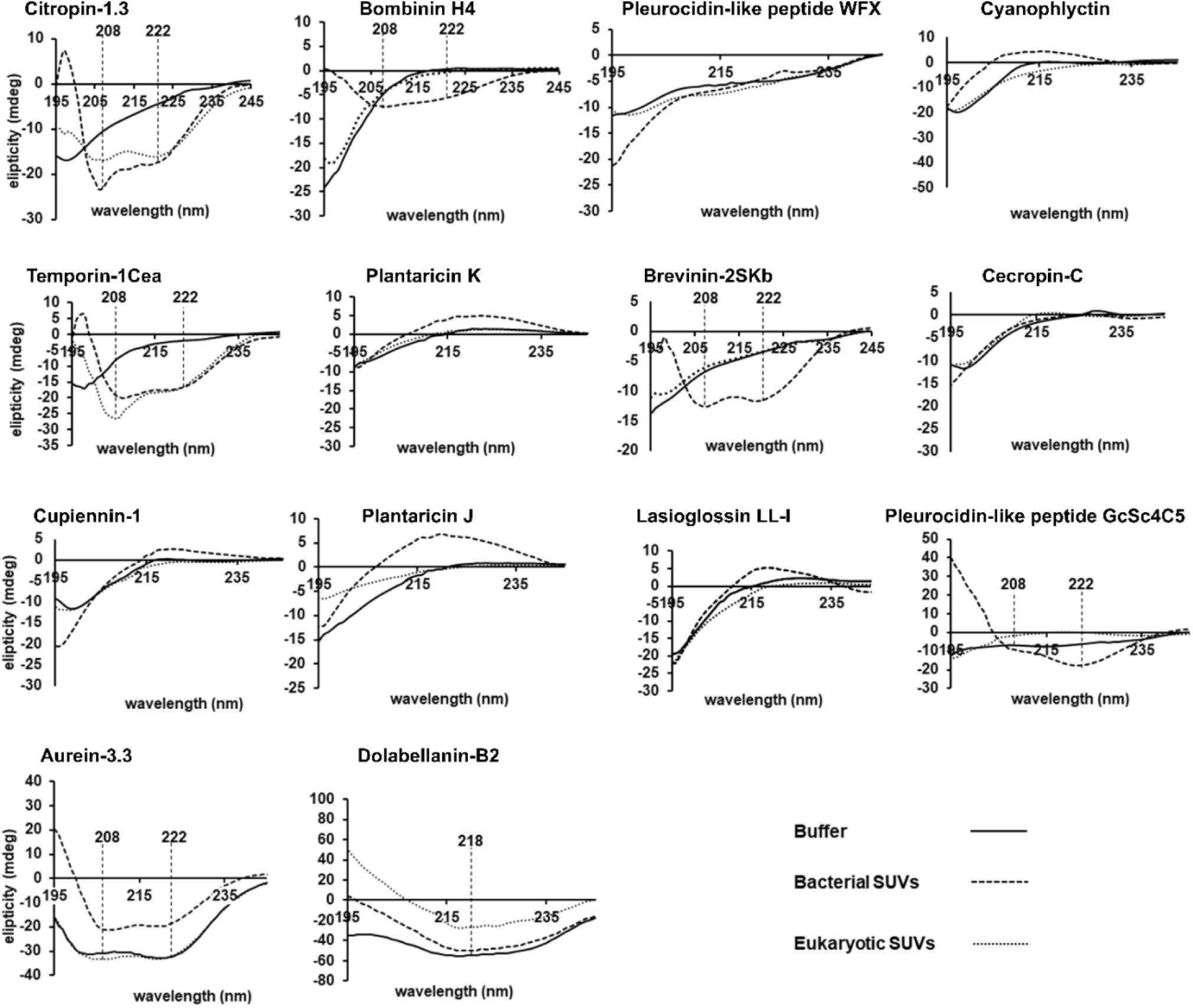
Secondary structure analyses of the ffAMPs in solution, as measured by circular dichroism. Solution CD spectra of the ffAMPs. The peptides were dissolved in buffer without (solid curve) or with bacterial membrane mimics (SUVs of DOPE;DOPG) (dashed curve), or eukaryotic membrane mimics (SUVs of DOPC:Chol:SM) (dotted curve). The main secondary structure indicated by the spectra is listed in Supplementary Table 6.

**Supplementary Figure 5.**
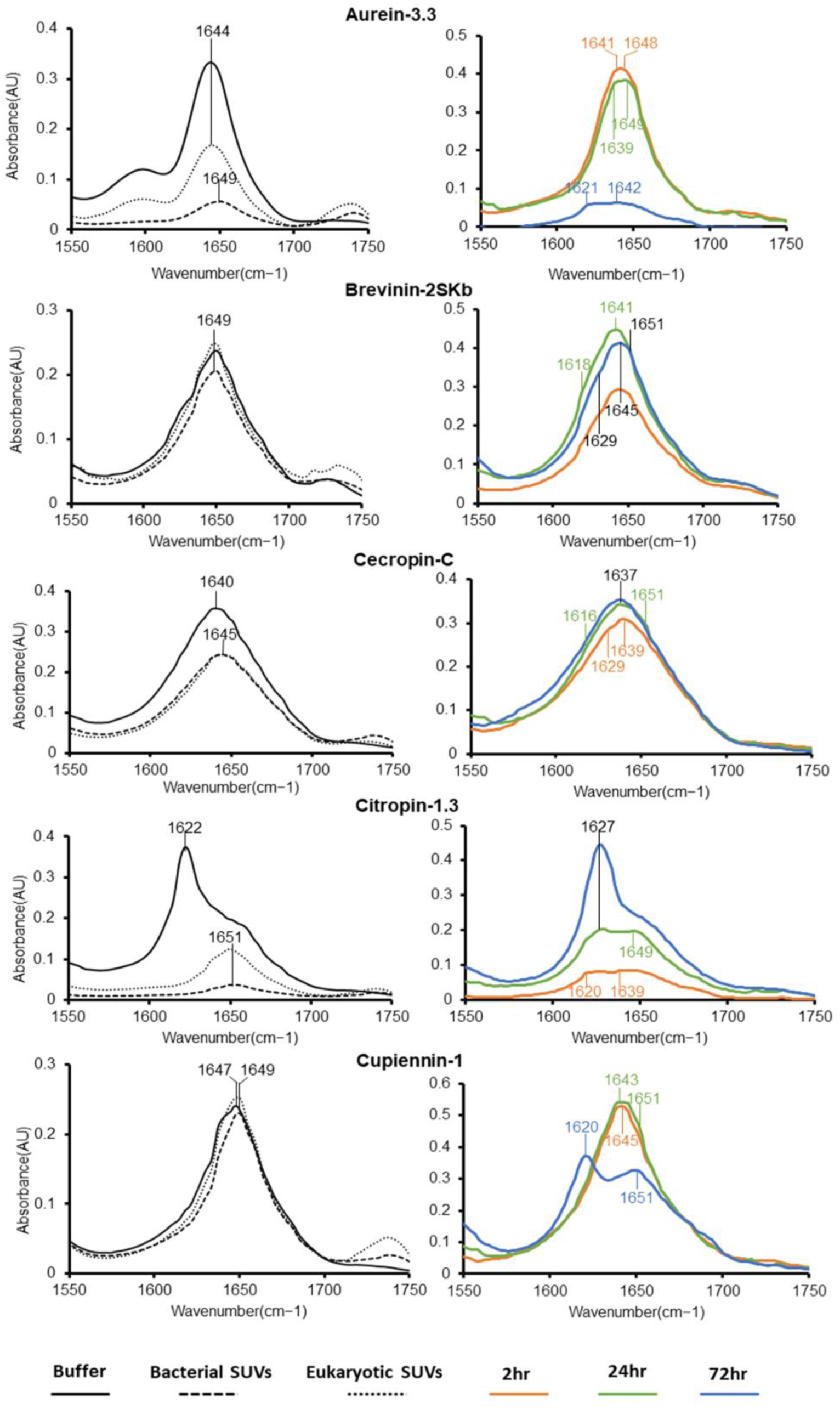
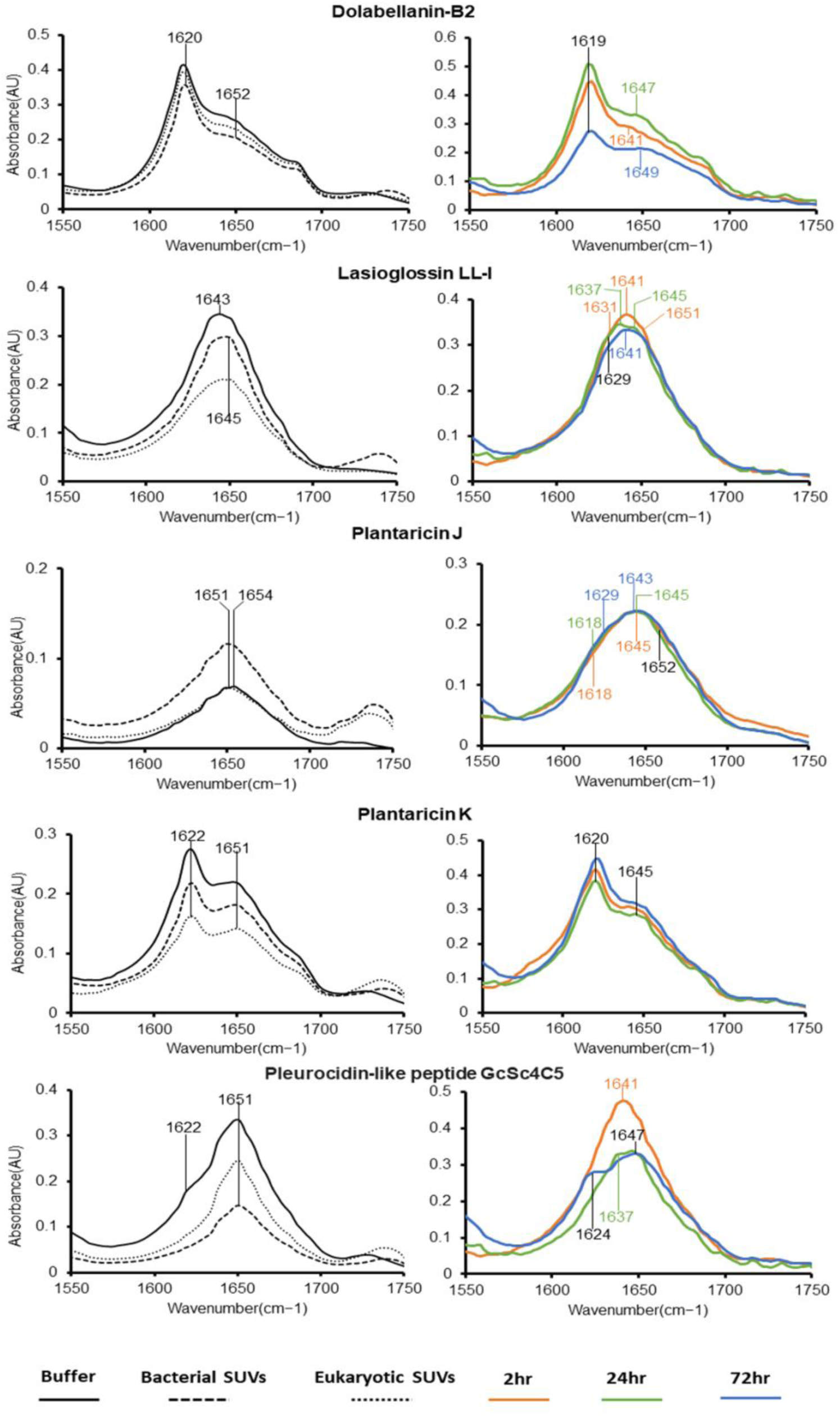
Secondary structure FTIR analyses of the ffAMPs in the solid/ fibrillar form. Attenuated total internal reflection Fourier transform infrared (ATR-FTIR) spectra of the amide I′ region of the ffAMPs under different conditions and over time. In the left panels, the spectra are of the peptides incubated for several hours in buffer (solid curve), or in the same buffer in the presence of SUVs composed of either DOPE:DOPG, mimicking a Gram-positive bacterial membrane (dashed curve), or DOPC:Chol:SM, mimicking an eukaryotic membrane (dotted curve). In the right panels, the spectra are of the peptides incubated in D2O for 2 h (orange curve), 24 h (green curve) or 72 h (blue curve). Peaks in the region of 1611–1630 cm^-1^ as well as ∼1685–1695 cm^-1^ are indicative of rigid cross-β fibrils, 1637–1645 cm^-1^ indicate disordered species, partially overlapping with peaks at 1630–1643 cm^-1^ correlating with small and disordered β-rich amyloid fibrils with absorbance which is typical of bent β-sheets in native proteins, and 1645–1662 cm^-1^ indicate the existence of α-helices ^17–20^, which overlap with random coil structures, especially for broad peaks ^21,22^ (both are therefore indicated in Table 1). The peaks are indicated based on the second derivative calculated by OPUS software.

